# A cingulate-hippocampal circuit mediates early depressive symptoms in Alzheimer’s disease

**DOI:** 10.1101/2023.02.07.527491

**Authors:** Yanbing Chen, Huimin Peng, Kai Zhuang, Wenting Xie, Chenli Li, Jin Xue, Meiqin Chen, Xiaoting Huang, Tingting Zou, Ya Wang, Dan Can, Huifang Li, Tifei Yuan, Jie Zhang

**Affiliations:** Fujian Provincial Key Laboratory of Neurodegenerative Disease and Aging Research, Institute of Neuroscience, Medical College, Xiamen University, Xiamen, Fujian, 361102, P.R.China.; College of Basic Medicine, Hebei Medical University, Shijiazhuang, Hebei, 050017, P.R.China.; Shanghai Mental Health Center, Shanghai Jiaotong University School of Medicine, Shanghai, 200000, P.R.China.

## Abstract

Depressive symptoms are prevalent and precedes cognitive decline in Alzheimer’s disease (AD), which worsen the clinical outcome and severely challenge the life quality of the patients. However, the neural circuitry mediating mood disturbance in AD remains elusive. Here we report that the glutamatergic projection from the anterior cingulate cortex (ACC) to the ventral hippocampal CA1 (vCA1) acts as the neural substrate of depressive symptoms in AD. We systemically mapped axonal projection in early stage of 5xFAD mice (3 months), which accompanies depressive-like behavior in these animals and identified reduced axonal projection from ACC to vCA1. This is in accordance with reduced axonal calcium signals detected with in vivo fiber photometry. Chemogenetic or optogenetic reversal of ACC-vCA1 circuitry activity efficiently ameliorated the depressive-like behaviors as well as cognitive impairment in 5xFAD mice. We further identified the synaptic molecules neuregulin-1 (Nrg1) as one vital signal regulating the synaptic glutamate transmission from the ACC to the vCA1. Collectively, these results indicated ACC-vCA1 as a critical pathway that regulates emotional states in Alzheimer’ disease. Targeting cingulate cortices with brain stimulation may treat depression in AD.

## Introduction

Alzheimer’s disease (AD) is characterized by irreversible neurodegeneration and amyloid plaque accumulation in the brain. Depression is highly prevalent in AD and severely worsens its clinical outcome, e.g., causing suicide (Even & Weintraub, 2010; D. H. Kim, Kim, Han, Jeon, & Han, 2020; Novais & Starkstein, 2015). However, in clinical practice currently it lacks effective treatment for depression in AD (D-AD). The available and first-line treatment approaches for major depressive disorder (MDD) (e.g., selective serotonin reuptake inhibitors (SSRIs)) showed low efficiency for depression symptoms in AD patients (Chan et al., 2020; Lee & Lyketsos, 2003). Therefore, it is critical to elucidate the potential neural mechanisms for depression in AD, in order to develop novel treatment approaches.

Recent neuroimaging findings reported decreased functional connectivity between ACC and the right occipital cortex and right lingual gyrus in D-AD patients, when compared to non-depressed AD subjects (nD-AD) (X. Liu et al., 2017). These findings were in accordance with other evidences supporting the involvement of ACC in major depression (Greicius et al., 2007; Mayberg et al., 2005; Nofzinger et al., 2005). Targeting ACC activity is also implicated in the rapid anti-depressant effects of ketamine (Alexander, Jelen, Mehta, & Young, 2021). In fact, ACC sends dense projection to hippocampus, which are important for emotion, pain and fear memory (Chen et al., 2019; Tao et al., 2019), and may contribute to its functioning in depression treatment. Notably, previous findings found that the projection from the ACC to ventral hippocampal CA1 (vCA1) regulates emotional memory generalization (Bian et al., 2019; Rolls, 2019). It is conceivable that the ACC-vCA1 pathway is involved in emotional and cognitive processing, and is implicated in depression in AD.

Here we employed a transgenic mice model of AD, the 5xFAD mice, carrying five familial AD mutations in human PS1 and mutant human APP, overproducing Aβ in the brain to assess the neural mechanism underlying the associated depressive like behaviors. We performed quantitative whole-brain of ACC-vHPC axonal pathways using viral-based anterograde tracing. The fiber photometry recording was employed to reveal that the activity was suppressed from ACC to vCA1 in early stage of AD mice relative to wile-type (WT) mice. Optogenetic and chemogenetic activation of ACC- vCA1 projection significantly ameliorate depressive-like behaviors and spatial memory deficits in 3 months- and 6 months-old of AD mice, respectively. Then we performed RNA-Sequencing combined with photostimulation to investigate the molecular mechanism regulating the process of depression-associated cognitive defects. Neuregulin-1(Nrg1) is the vital signaling molecule that modulated synaptic glutamate transmission in the ACC-vCA1 circuit. These results shall elucidate the potential involvement of ACC-vCA1 pathway in regulating depression-AD comorbidity and point out the relevant molecular target for therapeutic intervention.

## Results

### Reduced axonal projections and synaptic transmission of ACC-vCA1 circuit in 5xFAD mice

We firstly injected anterograde viral tracer AAV-CamkIIα-EYFP into ACC region (Figure 1A) and quantified the axonal projection in the whole brain (Figure 1B). Notably, the projections of ACC to several brain region are impaired (Figure 1B-C), including striatum (CPU), anterior part (PVA), thalamic reticular nucleus (TRN), basolateral amygdaloid nucleus (BLA), field CA1 of hippocampus (CA1), and dorsal raphe nucleus (DRN). We further carefully investigated the fact that the ACC projection terminals decreased dramatically in vCA1 (Figure 1D). We used a recombinant rabies virus (RV) retrograde monosynaptic tracing system to dissect the transsynaptic projection from ACC to vCA1. The Cre-dependent helper virus mixture (1:1) of AAV- EF1a-DIO-GT and AAV-EF1a-DIO-RVG was unilaterally injected into vCA1 of Camk2α-cre mice. Three weeks later, the RV-EnvA-DG-DsRed was injected into the vCA1 at the same site. The helper virus allows the RV spread one synapse retrogradely (Figure 1—figure supplement 1A-C), and DsRed signal was detected in ACC (Figure 1—figure supplement 1D). Double-immunofluorescent staining was used to identify the neurochemical properties of DsRed retrogradely labeled. Specifically, the majority of DsRed labeled neurons contained Camk2 (96%), while only 4% of DsRed labeled neurons were GAD67-positive in the ACC (Figure 1 — figure supplement 1E-G), suggesting that the ACC sends inputs to pyramidal cells but not interneurons in the vCA1.

**Figure 1.**
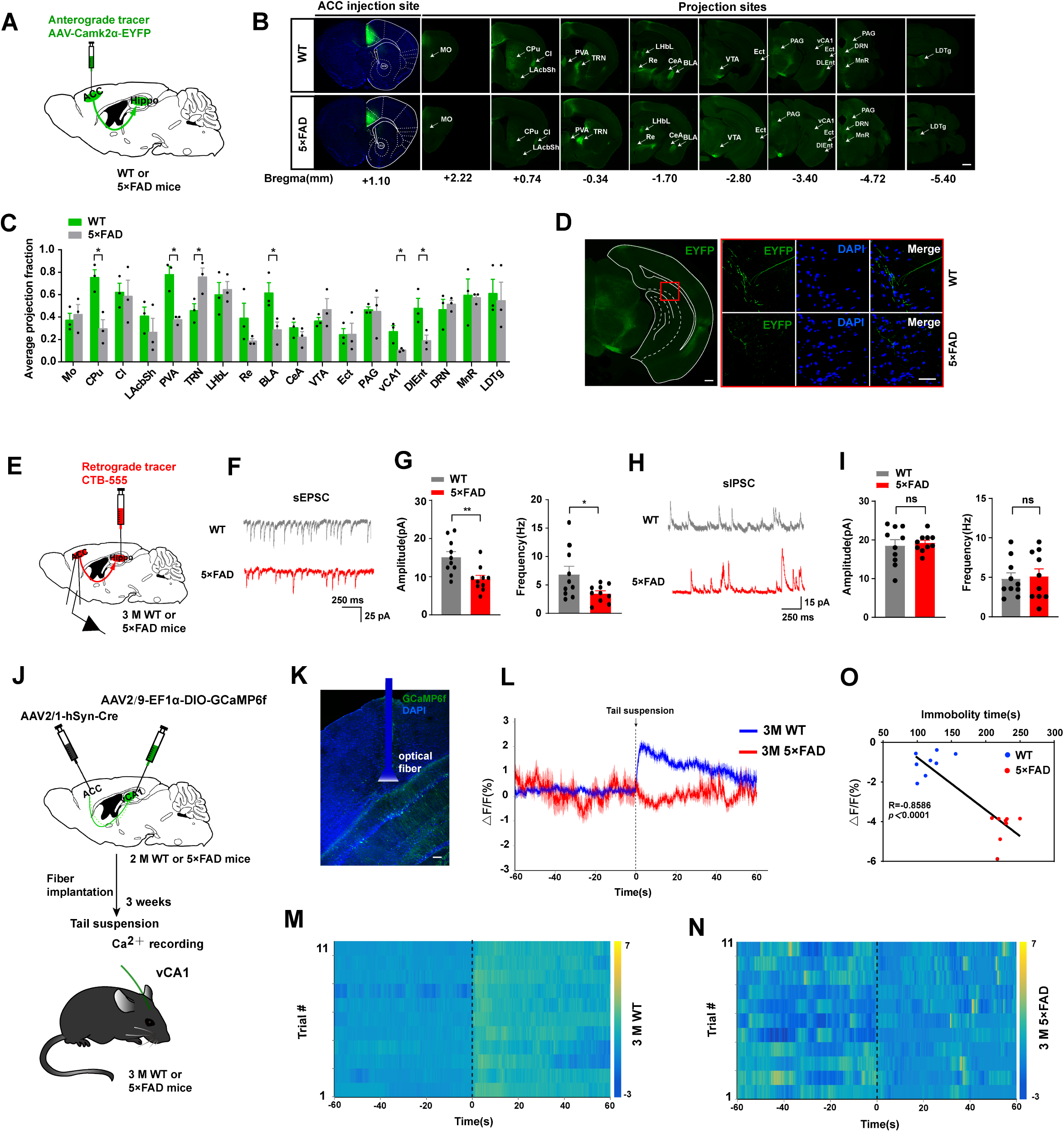
Glutamatergic ACC-vCA1 synaptic transmission was depressed in 3-months old 5xFAD mice. **(A)** The schematic diagram of the anterograde viral tracer AAV- Camk2α-EYFP was unilaterally injected into the ACC of 2 months-old 5xFAD mice or WT. (**B)** Series of coronal section from ACC pyramidal neurons in control and AD mice after injection of virus (scale bar, 500 μm). Mo, medial orbital cortex; CPU, striatum; Cl, claustrum; LAcbSh, lateral accumbens shell; PVA, paraventricular thalamic nucleus, anterior part; TRN, thalamic reticular nucleus; LHbL, lateral habenular nucleus, lateral part; Re, reuniens thalamic nucleus; BLA, basolateral amygdaloid nucleus; CeA, central amygdaloid nucleus; VTA, ventral tegmental area; Ect, ectorhinal cortex; PAG, periaqueductal gray; CA1, field CA1 of hippocampus; DlEnt, dorsolateral entorhinal cortex; DRN, dorsal raphe nucleus; MnR, median raphe nucleus; LDTg, laterodorsal tegmental nucleus. (**C)** Average fraction of ACC pyramidal neural axons in 18 brain regions. Projection fractions were normalized by the florescence density of the injection site. (WT, n = 3; 5xFAD, n = 3; unpaired *t* test; CPu, *t* = 4.283, *P* = 0.0128; PVA, *t* = 5.185, *P* = 0.0076; TRN, *t* = 2.962, *P* = 0.0415; BLA, *t* = 2.875, *P* = 0.0451; vCA1, *t* = 3.099, *P* = 0.0363; DlEnt, *t* = 2.860, *P* = 0.0459). **(D)** Representative image of the ventral hippocampal CA1 region in AD and WT mice (scale bar 500 μm and 20 μm in the red rectangle). **(E)** Schematic showing the injection of Cholera toxin subunit B (recombinant), Alexa Fluor-555 conjugate (CTB-555) into the vCA1 and whole-cell patch recording of ACC neurons. Representative traces (**F**) and average data showing that the frequency and amplitude (**G**) of sEPSC of CTB-555^+^ neurons are significantly weakened in 5xFAD mice compared with control mice (amplitude: *t* = 4.652, *P* = 0.022; frequency: *t* = 2.245, *P* = 0.0376; mann–Whitney test; n = 10 cells from three mice per genotype). Representative traces (**H**) and average data showing that the frequency and amplitude (**I**) of sIPSC of CTB-555^+^ neurons are no significantly changes between 5xFAD mice and control mice (amplitude: t = 0.4053, *P* = 0.6900; frequency: t = 0.2416, *P* = 0.8118; mann–Whitney test; n =10 cells from three mice per genotype). (**J)** Schematic of AAV2/1-hSyn-Cre and AAV2/9-EF1α-DIO-GCaMP6f injection into the ACC and vCA1, respectively, and Ca^2+^ recording in the TST. (**K)** Representative image of GCaMP6f expressing cells and optic fiber track in the vCA1 (scale bar 100 μm). The mean (**L**) and heatmaps (**M**, **N**) show that Ca^2+^ signals rapidly decreased in 3 months- old 5xFAD mice compared with control mice when subjected to tail suspension. The colored bars of (**N**) and (**O**) indicate ΔF/F (%). (**O)** Relationship between Ca^2+^ signals in vCA1 and immobility time in FST (Pearson’s correlation, *R* = −0.8586, *P* < 0.0001; n = 8 mice per group). Data represent mean ± SEM, ns: not significant, \**P* < 0.05; \*\**P* < 0.01; \*\*\**P* < 0.001. Figure 1**—source data 1.** Source data indicating GCaMP6f fluorescent signals align to tail suspension test. Figure 1—figure supplement 1. The direct glutamatergic monosynaptic projection from ACC to vCA1.

Then we employed ex vivo brain slice recording to measure the synaptic transmission onto the ACC-vCA1 projecting neurons in early stage of 5xFAD mice. We used Cholera toxin subunit B (recombinant), Alexa Fluor-555 conjugate (CTB-555) to retrogradely label the ACC-vCA1 projection neurons and then patch these CTB-555- positive neurons in the ACC of brain slices in electrophysiological study (Figure 1E). In the voltage clamp, both the amplitude and frequency of spontaneous excitatory postsynaptic currents (sEPSC) (Figure 1F-G) in 3-month-old 5xFAD were decreased compared with control. However, there is no changes in both the frequency and amplitudes of spontaneous inhibitory postsynaptic currents (sIPSC) (Figure 1H-I).

To further investigate the neural activity in ACC-vCA1 pathway, we performed axonal fiber photometry recording of Gcamp6f-expressing ACC to vCA1 pathway activity in mice subjected to TST on 3-month-old 5xFAD mice. AAV serotype 1 expressing recombinase Cre (Zingg et al., 2017) was as an anterograde transsynaptic tracer (AAV2/1-hSyn-Cre) and AAV2/9-EF1α-DIO-GCaMP6f were injected into the ACC and vCA1, respectively, and an optic fiber was implanted into the vCA1 (Figure 1J, K). Notably, the Ca^2+^ signals were much lower in 5xFAD mice, when compared to WT mice upon tail suspension test (Figure 1L-N); there was a strong negative correlation between the level Ca^2+^ signals and immobility time in the tail suspension test (TST) (Figure 1O). These data indicated reduced excitatory synaptic transmission in ACC-vCA1 pathway, accompanying the depressive like behaviors in early stage of 5xFAD mice.

### Chemogenetic activation of ACC-vCA1 projections attenuate the depressive behaviors and the spatial memory deficit in 5xFAD mice

Next, we assessed the effects of activating ACC-vCA1 pathway on the depressive-like behavior of 5xFAD mice. The DREADDs (Gq-coupled designer receptor, hM3D) and its ligand CNO were used to selectively manipulate the activity of ACC-vCA1 circuit, as shown in Figure 2. These mice were then subjected to related behaviors tests at 3 months old and 6 months old, respectively (Figure 2B). Notably, the chemogenetic activation of the ACC-vCA1 pathway by CNO significantly reversed the depressive-like behaviors in 5xFAD mice by TST, FST and SCP tests (Figure 2C-D). In Morris water maze (MWM), the learning ability was increased by activation ACC-vCA1 projection on 6-month-old 5xFAD mice (Figure 2E). In addition, during the probe trial, the chemogenetic activation of ACC-vCA1 inputs robustly increased the time in target quadrant, entries into the platform location and required a shorter period to travel from the entry point to the target zone without changing motor ability in 5xFAD mice (Figure 2F-I). However, no differences were found by inhibition of such neuronal pathway (Figure 2 — figure supplement 1). The efficacy of hM3D-mediated activation was confirmed with FOS immunofluorescence staining (Figure 2J-K) and slice recording (Figure 2L-N). There were numerous mCherry-expressing neurons in the vCA1 and these neurons were the ones receiving monosynaptic inputs from the ACC (Figure 2J and 2O-P). Although no difference in plaque number was detected in vCA1 after CNO-injection intraperitoneally (Figure 2—figure supplement 2), activation of ACC-vCA1 could restore spatial memory dysfunction in 5xFAD. Collectively, these results suggest that activating ACC-vCA1^Camk2^ pathway ameliorate depression in early stage of AD mice and spatial memory deterioration of 5xFAD mice.

**Figure 2.**
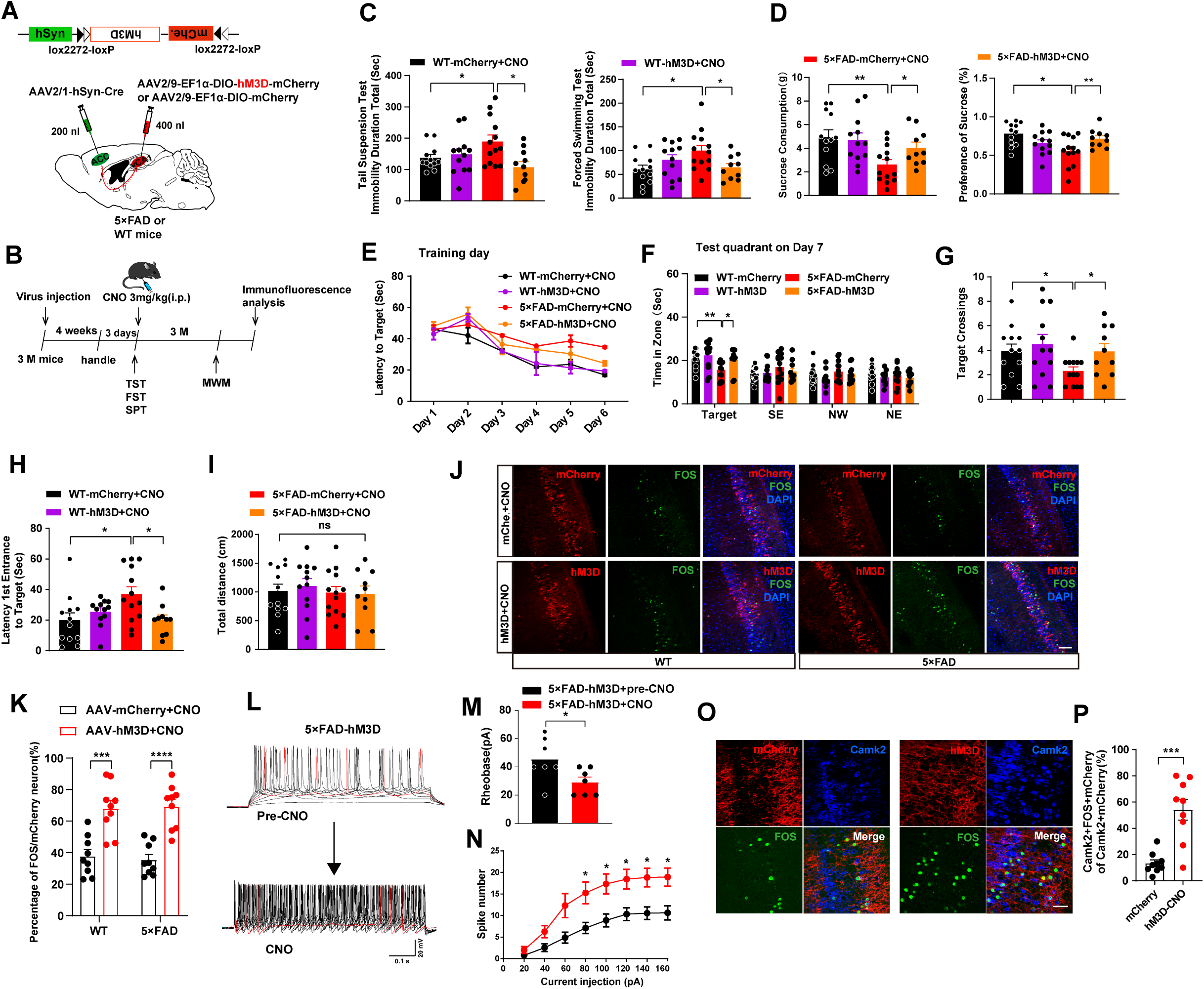
Chemogenetic activation of ACC-vCA1 projections led to antidepression effect in early stage of AD mice and rescue subsequent spatial memory impairment. Schematic diagram (**A**) and behavioral experimental timeline (**B**) of virus injection and optical fiber implant in 2 months-old and age-matched WT and 5xFAD mice. AAV2/1- hSyn-Cre is injected into the ACC, and AAV2/9-DIO-hM3D-mCherry is injected into the ipsilateral vCA1. The effect of ACC-vCA1 pathway chemogenetic activation on early depressive-behaviors in 3 months-old 5xFAD mice in TST, FST (**C**) and SPT (**D**). n = 12, 12, 13 and 10 WT-mCherry+CNO, WT+hM3D+CNO, 5xFAD+mCherry+CNO and 5xFAD+hM3D+CNO group, respectively. During Morris water maze tests, WT+mCherry, WT+hM3D, 5xFAD+mCherry and 5xFAD+hM3D mice receiving CNO injections were analyzed for escape latency during a 6-day training period (**E**). On the next day, mice were analyzed for time spent in the target zone and other quadrants (SE, NW and NE) (**F**), number of target crossings (**G**) and time required from entrance to the target platform (**H**) without changes in total distance (**I**). n = 12/12/13/10 mice per group. (**J)** Immunofluorescent images showing viral expression, FOS expression and co-labeled neurons in vCA1 of both WT and 5xFAD mice. (**K)** Proportion of FOS- positive cells that co-express mCherry. n = 9 slices from 4 mice per group (scale bar 50 μm). **(L)** Current–voltage relationship of a representative vCA1 mCherry-labeled projective neuron recorded before, and after CNO perfusion (5 μM). Raw traces show individual voltage responses to a series of, 600 ms current pulses from 0 to 200 pA in 20 pA steps. Red traces indicate number of 200 pA current-induced action potentials. (**M)** The rheobase current was decreased by CNO (5 μM) administration. Unpaired t-test. (**N)** Number of induced APs at different current steps, n = 7 neurons from 3 mice. (**O**) Immunofluorescent images (scale bar 25 μm) and column diagram (**P**) of mCherry/FOS/Camk2 triple-labeled staining in vCA1 of 5xFAD-mCherry+CNO and 5xFAD-hM3D+CNO group. unpaired t-test, n = 9 slices from 5 mice per group. Data represent mean ± SEM. ns: not significant, \**P* < 0.05; \*\**P* < 0.01; \*\*\**P* < 0.001, \*\*\*\**P* < 0.0001. Figure 2**—source data 1.** Source data indicating chemogenetic activation of ACC- vCA1 neural circuit on some control behaviors. Figure 2**—figure supplement 1.** No significant behavioral changes were found by chemogenetic inhibition of ACC-vCA1 pathway. Figure 2**—figure supplement 2.** Chemogenetic activation rescues ACC-vCA1 without changing Aβ level in vCA1.

### Activation of the ACC-vCA1 inputs enhances synaptic transmission

Synaptic dysfunction contributes to alternations in learning and memory. One of the main forms of synaptic plasticity found at the excitatory synapses is the long-term potentiation (LTP), which requires the activation of postsynaptic glutamate receptors and is altered in neurodegenerative diseases (Brzdak, Nowak, Wiera, & Mozrzymas, 2017). Here we observed a substantial reduction in high frequency stimulation (HFS)- induced LTP in 5xFAD mice, while chemogenetic activation of ACC-projecting vCA1 glutamate neurons in 5xFAD mice augments this LTP (Figure 3A-C). Golgi staining indicates that the spine generation significantly decreased in AD mice compared with the WT, whereas activation of ACC-vCA1 by hM3D-chemogenetical treatment restored the decrease without changes in inhibition of the circuit (Figure 3D-E), suggesting a weakened synaptic plasticity of ACC-vCA1 neural circuit in AD mice. The decreases of synaptic protein such as GluR1, GluR2 and PSD95 in 5xFAD mice were also rescued by chemogenetic activation of ACC-vCA1 (Figure 3F-G). However, no significant change was observed with the inhibitory chemogenetic manipulations (Figure 3F-G). The activation of ACC-vCA1 does not affect the APP expression (Figure 3—figure supplement 1).

**Figure 3.**
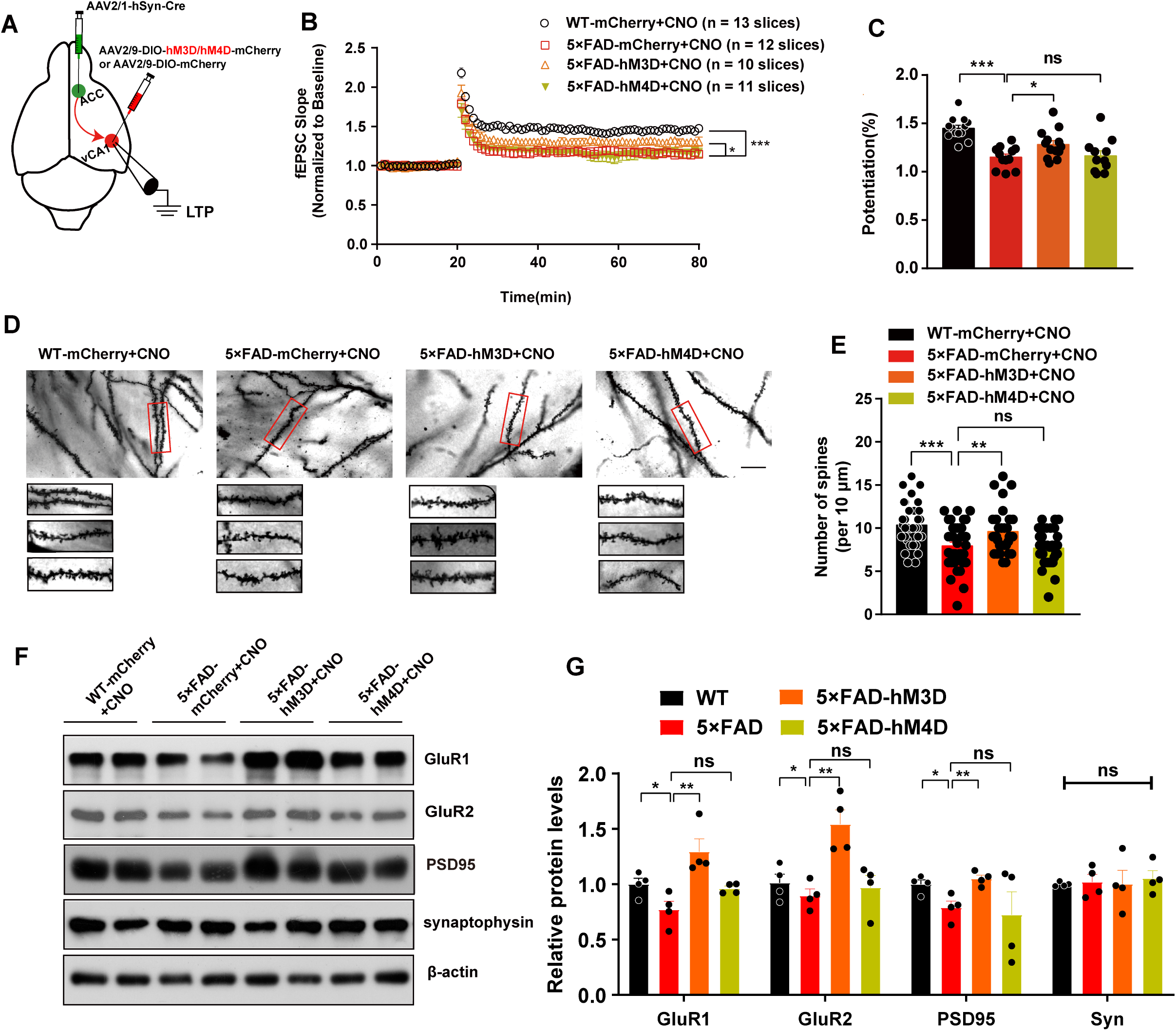
Postsynaptic plasticity is restored via activation but not inhibition of the ACC-vCA1 inputs in AD mice. **(A)** Schematic diagram showing the injection of the virus and the placement of recording electrodes in vCA1. AAV2/1-hSyn-Cre is injected into the ACC, and AAV2/9-DIO-hM3D-mCherry or AAV2/9-DIO-hM4D-mCherry is injected into the ipsilateral vCA1. (**B)** Time course of the averaged fEPSP slope and amplitude of the LTP recordings in each group. fEPSP potentiation was quantified during the last 10 min of recording, and significance was determined by repeated- measures ANOVA with Bonferroni’s post hoc analysis. **(C)** Column diagram showing that average slope of fEPSP in the 6 months-old 5xFAD mice is decreased compared to age-matched control mice, chemogenetic activation but not inhibitation of the ACC- vCA1 neural pathway in 5xFAD mice rescues those effect. n = 13/12/10/11 slices from 5 mice per groups. (**D**) Representative Golgi staining images (scale bar 10 μm) and quantification (**E**) of dendritic spines from vCA1 in four groups. n = 36 dendritic segments from 3 mice per group. (**F)** Representative images of western blot. (**G)** Quantification of GluR1, GluR2, PSD95 and synaptophysin in vCA1 after chemogenetic activation or inhibition of ACC-vCA1 pathway. n = 4 mice per group. Data represent the mean ± SEM. One-way ANOVA analysis. ns: not significant, \**P* < 0.05, \*\*\**P* < 0.001. Figure 3**—source data 1.** (related to Figure 3F and G) Immunoblot and quantification of GluR1, GluR2, PSD 95 and synaptophysin in hippocampal CA1 after chemogenetic activation or inhibition of the ACC-vCA1 circuit. Figure 3**—figure supplement 1.** The activation of ACC-vCA1 does not affect the APP expression.

### Optogenetic-stimulation of the ACC-vCA1 circuit ameliorates depression-associated memory impairment in AD mice

We next employed another strategy to activate ACC-vCA1 by optogenetic activation of this projection. In brief, mice were given stereotaxic injections of an anterograde recombinant AAV receptors virus expressing Cre recombinase controlled by hSyn promotor (AAV2/1-hSyn-Cre), following injection an adeno-associated virus with Cre dependent constructs encoding ChR2 (AAV-EF1α-DIO-ChR2-mCherry) into the ACC and vCA1, respectively. Optic fibers were inserted into the vCA1 before behavioral testing (Figure 4A-B). To assess the functional consequence of increased neuronal firing in ACC-vCA1 projections, the behavioral effects of optogenetic-induced phasic firing (20Hz, 5ms), in ChR_2_-expressing vCA1 neurons were evaluated during depression-behavioral and MWM testing. Notably, blue light stimulation significantly activated the vCA1 excitatory neurons innervated by ACC (Figure 4C-D) and attenuates the depressive behaviors tested by SPC, TST and FST (Figure 4E-H). The learning and memory defects were also rescued by targeted photostimulation of ACC- vCA1 input in 5xFAD mice (Figure 4I-L) without effect on moving ability (Figure 4M).

**Figure 4.**
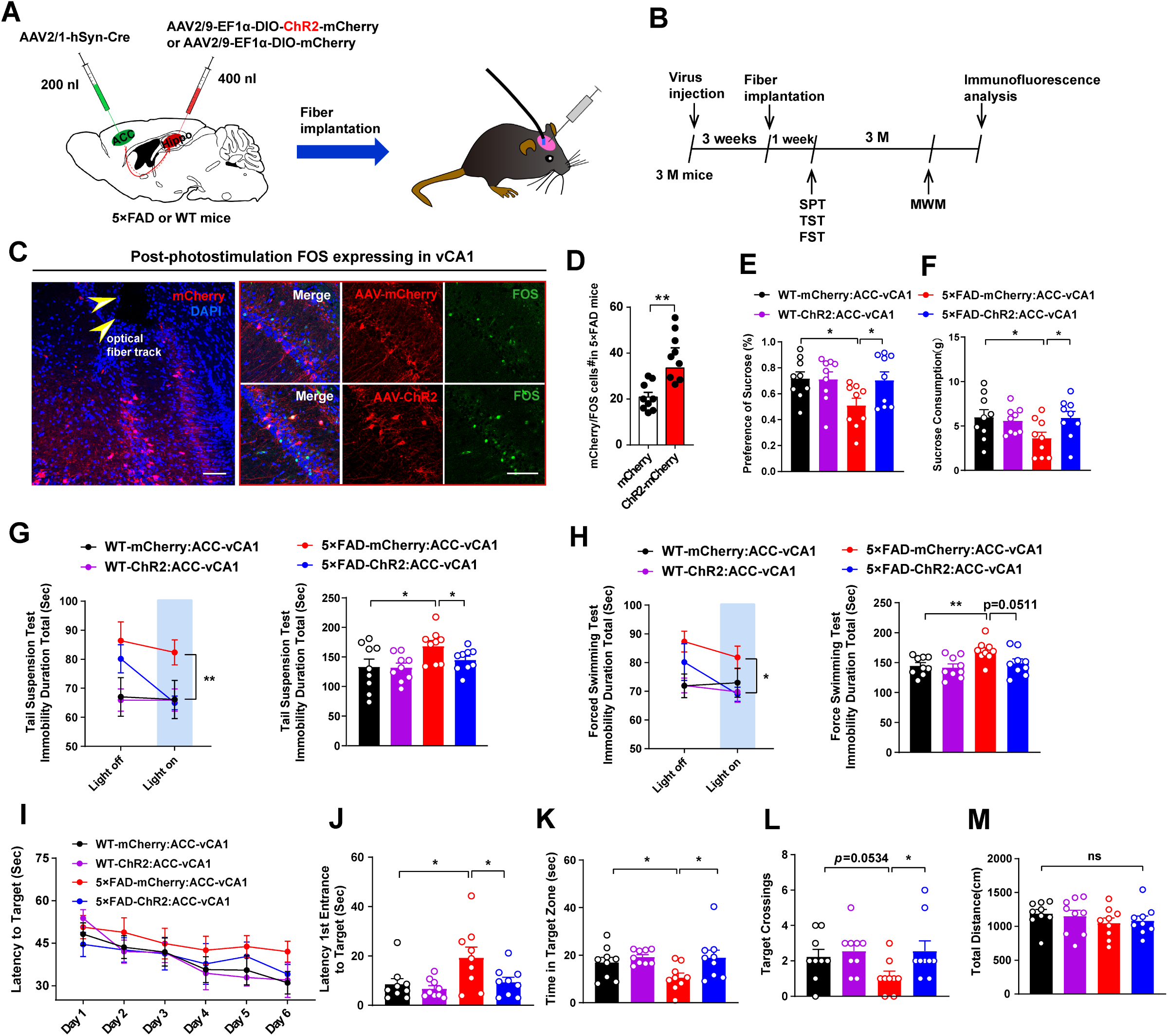
Photostimulation of the ACC-vCA1 circuit ameliorates early depressive-like behaviors and subsequent depression-associated memory impairment in AD mice. (**A**) Schematic diagram and behavioral experimental timeline (**B**) of virus injection and optical fiber implant in 2 months-old and age-matched WT and 5xFAD mice. (**C)** Immunofluorescent images showing virus injection, optical fiber placement and FOS^+^ neurons in the vCA1 of the photo-stimulated mice. Yellow arrowheads indicate optical fiber track (scale bar 50 μm and 25 μm in the red rectangle). (**D)** Column diagram showing the number of FOS^+^/mCherry double-labeled cells are increased by photostimulation of ChR_2_-expressing ACC-vCA1-projection neurons in 5xFAD mice. unpaired t-test, n = 9 slices from 5 mice per groups. The effect of ACC-vCA1 pathway photoactivation on early depressive-behaviors in 3 months-old 5xFAD mice in SPT (**E, F**), TST (**G**) and FST (**H**). The immobility time in TST (**G, left**) and FST (**H, left**) is measured during the lights off and on (two sessions with 3 min for each session with photostimulation. (**I-L)** During Morris water maze tests, WT-mCherry mice, WT-ChR_2_ mice, 5xFAD-mCherry, and 5xFAD-ChR_2_ mice were analyzed for escape latency during a 6-day training period (**I**). On the next day, mice were analyzed for time required from entrance to the target platform (**J**), target zone (**K**), and number of target crossings (**L**). **(M)** Total distance during the training did not differ among four groups. Data represent mean ± SEM. One-way ANOVA analysis, n = 9 mice per groups. ns: not significant, \**P* < 0.05; \*\**P* < 0.01. Figure 4 **— source data 1.** Source data indicating behavioral performances of optogenetic modulation of ACC-vCA1 circuit in 3-month-old WT and AD mice.

### Nrg1 is required for the regulation of emotional states and spatial memory deficit

At early stage of AD mice (3-month-old), dramatically amyloid plaque deposition was observed in ACC of AD mice compared with control (Figure 5A). We collected ACC from AD and control mice brain and subjected them for RNA-sequence (RNA-seq) analysis (Figure 5B). The results showed that 835 genes increased and 179 genes decreased in ACC from AD mice brain, when compared to control (Figure 5C). Kyoto Encyclopedia of Genes and Genomes (KEGG) pathway enrichment analyses and Gene Ontology (GO) were performed for the differential screens of the expression of genes (DEGs) (Figure 5—figure supplement 1A and B). Among the pathway clustered in KEGG analysis, the “neurodegenerative disease” list at the top; GO term of cellular component (CC) analysis also indicates that genes in synapse is dramatically affected (Figure 5D-E). Considering the vital role of synaptic function in depression and cognition regulation, we combine and analysis the overlap DEGs both in “neurodegenerative disease” and “synapse” (Figure 5F-G). In these overlapped genes, Neuregulin-1 (Nrg1) draws our attention. Nrg1 is highly involved in presynaptic vesicular transporting and exocytosis which mediates synaptic transmission (Chaudhury et al., 2003). Geng et al. showed that Nrg1 increased the frequency of sIPSC in hippocampus by in vitro slice recordings and its modulation of presynaptic release (Geng et al., 2016; Ozaki, Itoh, Miyakawa, Kishida, & Hashikawa, 2004). We found that the mRNA and protein levels of Nrg1 increased significantly in ACC of 3- month-old AD mice brain compared with control (Figure 5H). Immunofluorescent doubled staining further confirmed increased Nrg1 expression in pyramidal cells in ACC (Figure 6A-B and Figure 6—figure supplement 1).

**Figure 5.**
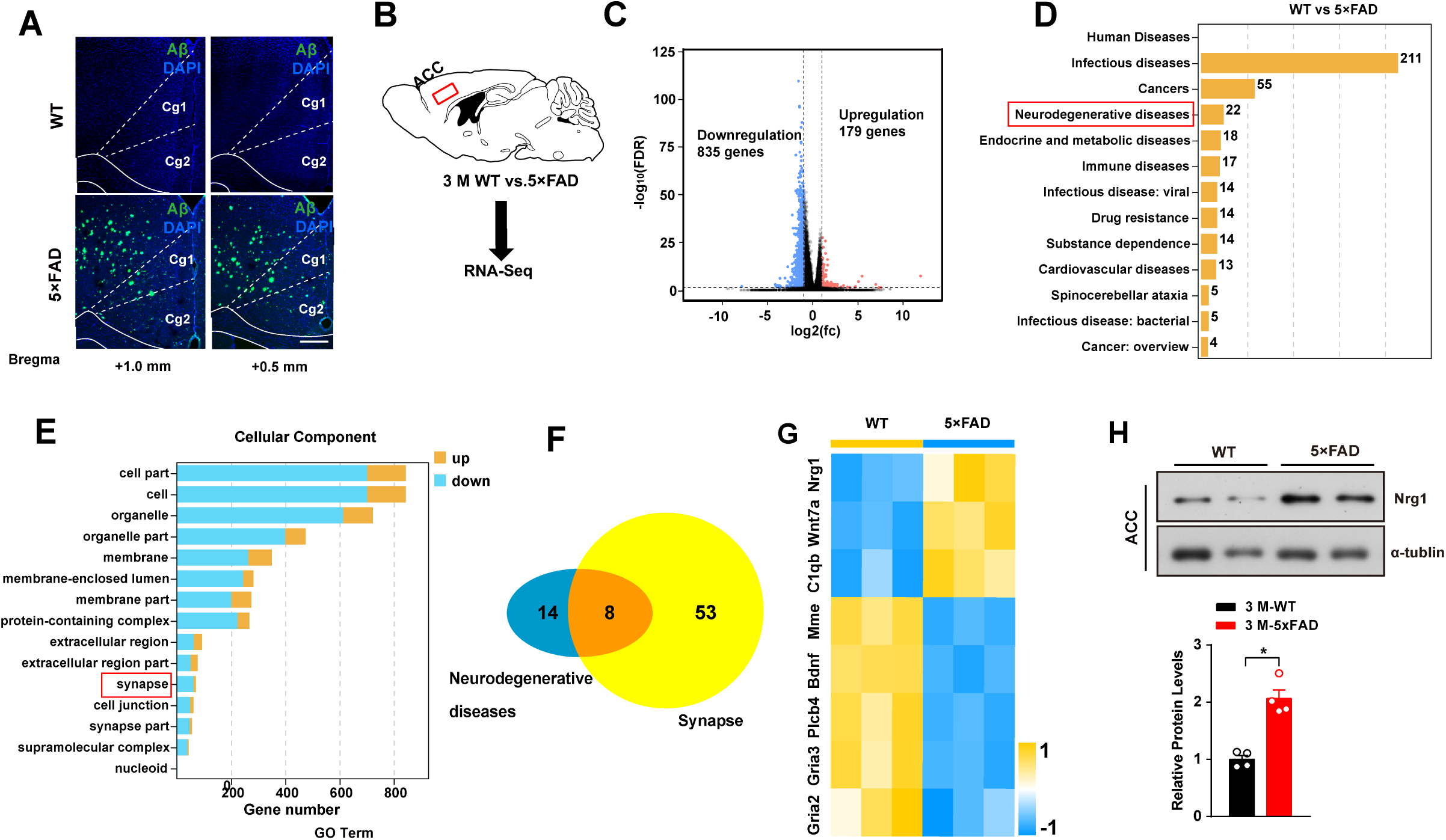
Identification of the key molecules involved in depression in early AD mice. **(A)** Representative image of Aβ pathology in the ACC by MOAB-2 staining (scale bar 100 μm). (**B)** Schematic showing the RNA sequence procedure. **(C)** Volcano plot showing changes in the expression of transcripts in the ACC between 3 months-old WT and 5xFAD mice. The y axis corresponds to the mean expression level of the log_10_ false discovery rate (FDR), and the x axis displays the log_2_ fold change. Red dots represent upregulated transcripts (*P* < 0.05, FDR *q* < 0.05) and blue dots represent downregulated transcripts (*P* < 0.05, FDR *q* < 0.05) between the WT and AD groups. The dashed line parallel to the x axis indicates a raw *q* value = 0.05. The dashed line parallel to the y axis indicates a raw FC = 1.5. (**D)** Differential expressing genes (DEGs) were clustered by Kyoto Encyclopedia of Genes and Genomes (KEGG) analysis, and the human disease pathways are shown. Red rectangle indicates the total of 22 DEGs in “neurodegenerative disease” KEGG pathway. (**E)** DEGs were clustered by Gene Ontology (GO) analysis and term of cellular component (CC) is shown. Red rectangle indicates the total of 61 DEGs in “synapse” GO term of cellular component (CC). (**F)** Venn diagrams showing the number of overlapping genes in “neurodegenerative disease” KEGG pathway and in “synapse” GO term of cellular component (CC). (**G**) Heatmap illustrating the expression levels of different signaling molecules related to synaptic transmission in the ACC of mice. Different groups are indicated on the y axis (yellow, WT; blue, 5xFAD). The x axis shows different signaling molecules, including Nrg1. (**H)** Samples (top) and column diagram (bottom) showing the western blotting results. The expression levels of the Nrg1 proteins are increased in the AD groups. Data represent mean ± SEM. unpaired t-test. n = 4 mice per group. \*\*\**P* < 0.001. Figure 5**-source data 1.** Source data indicating the dentification of Nrg1 involved in depression in early AD mice by RNA sequencing. Figure 5**—figure supplement 1.** Differential expression genes-GO/KEGG analysis.

**Figure 6.**
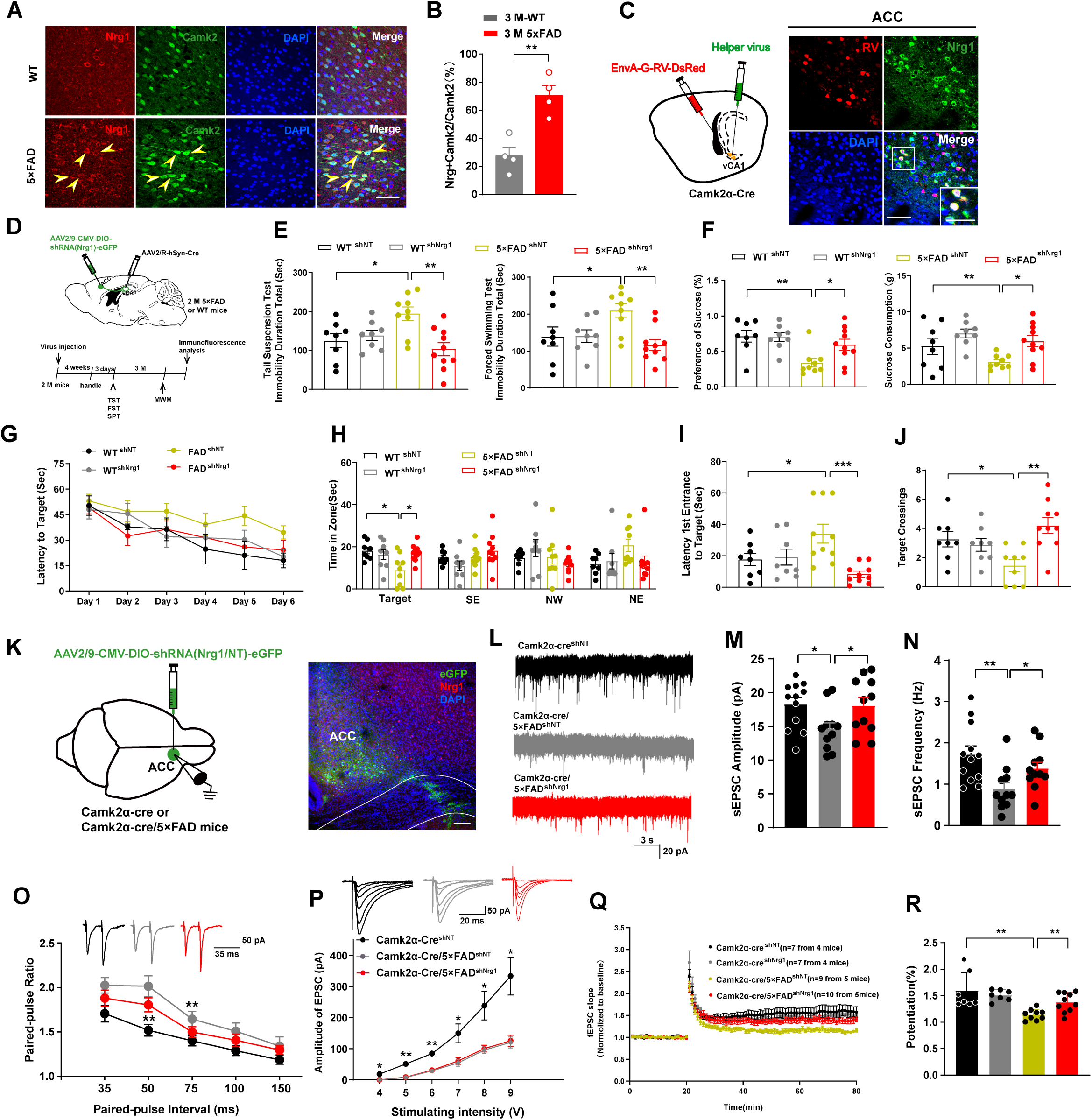
Downregulation of Nrg1 rescues early depressive-like behavior and cognitive deficit in AD mice. (A, B) Representative figures (**A**) (scale bar 20 μm) column diagram **(B)** showing the expression of Nrg1 in the Camk2 containing pyramidal cells in the ACC is much higher in 5xFAD mice than in WT mice. unpaired t test, t = 7.684, \*\**P* < 0.01, n = 4 mice per group. Yellow arrowheads indicate Nrg1 and Camk2 doubled- positive neurons. (**C)** Left: Schematic showing the injection of retrograde monosynaptic tracing in Camk2α-Cre mice: the helper viruses AAV-EF1α-DIO-GT and AAV-EF1α-DIO-RV-G (1:1, 200 nl, green) was injected into vCA1, and then rabies virus EnvA-pseudotyped RV-DG-DsRed (red) was also injected into vCA1. Right: The representative images of RV-DsRed co-labeling neurons in the ACC (scale bar 20 μm). White rectangle indicates RV-DsRed/Nrg1 double-labeled neurons (scale bar 5 μm). (**D)** Schematic diagram and experimental timeline showing the injection of the virus. AAV2/R-hSyn-Cre is injected into the vCA1, and AAV2/9-DIO-shRNA (Nrg1/NT)- eGFP is injected into the ACC, respectively. **(E)** Column diagram illustrating the immobility duration in WT + shRNA (NT), WT + shRNA (Nrg1), 5xFAD + shRNA (NT) and 5xFAD + shRNA (Nrg1) mice by TST (left) and FST (right). Immobility time in 5xFAD+shRNA (Nrg1) mice are decreased compared with those of scramble virus- injected mice. unpaired t-test, n = 8/8/9/10 mice per group. **(F)** Column diagram illustrating the sucrose preference and consumption of different groups by SPT and SCT. (**G-J)** During Morris water maze tests, WT + shRNA (NT), WT + shRNA (Nrg1), 5xFAD + shRNA (NT) and 5xFAD + shRNA (Nrg1) mice were analyzed for escape latency during a 6-day training period (**G**) On the next day, mice were analyzed for time spent in the target zone and other quadrants (SE, NW and NE) (**H**), time required from entrance to the target platform (I) and number of target crossings (**J**). (**K)** Schematic diagram and representative image showing the injection of the virus. AAV2/9-DIO- shRNA (Nrg1/NT)-eGFP is bilaterally injected into ACC of the Camk2α-Cre and Camk2α-Cre/5xFAD mice (scale bar 100 μm). **(L)** Representative sEPSC recorded in eGFP^+^ neurons of the ACC from Camk2α-Cre + shRNA (NT) (top), Camk2α- Cre/5xFAD+shRNA (NT) (middle) and Camk2α-Cre/5xFAD+ shRNA (Nrg1) (bottom) mice holding at -60 mV. (**M, N)** Summary plots showing the amplitude (**M**) and frequency (**N**) of sEPSC of was decreased in Camk2α-Cre/5xFAD+shRNA (NT) mice (n=11 neurons from 4 mice) compared to Camk2α-Cre + shRNA (NT) mice (n = 12 neurons from 4 mice) (unpaired t-test, amplitude: *t* = 2.578, *P* = 0.0175; frequency: *t* = 3.186, *P* = 0.0044). The amplitude (unpaired t-test, t = 2.187, *P* = 0.0408) and frequency (unpaired t-test, t = 2.246, *P* = 0.0361) were significantly potentiated in Camk2α- Cre/5xFAD+shRNA (Nrg1) mice (n = 11 neurons from 4 mice) compared to Camk2α- Cre/5xFAD+shRNA (NT) mice (n = 11 neurons from 4 mice). (**O)** Paired-pulse facilitation ratios were obtained from whole-cell recordings in ACC eGFP^+^ neurons. n = 12 neurons from 3 mice per group, one way ANOVA followed with Dunnett’s test. (**P)** The slope of the input–output curve reflecting as basal synaptic transmission is steeper in Camk2α-Cre + shRNA (NT) but not in Camk2α-Cre/5xFAD + shRNA (Nrg1) mice than that in the Camk2α-Cre/5xFAD + shRNA (NT) control mice. The ability of basal synaptic transmission is restored in Camk2α-Cre/5xFAD + shRNA (Nrg1) mice. n = 12 neurons from 3 mice per group. (**Q, R)** LTP recordings in vCA1 showing that fEPSP potentiation is enhanced by knocked down Nrg1-expressing in ACC pyramidal neurons in 5xFAD mice (Camk2α-Cre + shRNA (NT) versus Camk2α-Cre/5xFAD + shRNA (NT), *P* = 0.023; Camk2α-Cre/5xFAD+shRNA (NT) versus Camk2α- Cre/5xFAD + shRNA (Nrg1), *P* = 0.036; repeated-measures ANOVA with Bonferroni’s post hoc analysis). Data represent mean ± SEM. \**P* < 0.05; \*\**P* < 0.01. Figure 6**—source data 1.** Source data indicating presynaptic downregulation of Nrg1 relieved depressive-like behaviors and spatial learning and memory deficits in AD mice. Figure 6**—figure supplement 1.** The co-existence of Nrg1 with GAD67 in ACC. Figure 6**—figure supplement 2.** Nrg1 was knocked down efficiently by AAV-sh-Nrg1 microinjection into the ACC.

To investigate whether the Nrg1 expressed neuron mediates the ACC-vCA1 projection, the EnvA-G-Retro-virous was injected into vCA1, notably, the retrogradely labeled neurons (DsRed) from the vCA1 co-localized with Nrg1 signals in ACC (Figure 6C and Figure 1—figure supplement 1A-D). Since Nrg1 mostly locates and plays function in pre-synapse, we knockdown the presynaptic Nrg1 in ACC-vCA1 projection.

The knockdown virus (AAV2/9-DIO-shNrg1) was first injected into the ACC region, then AAV2/R-Cre virus was injected into the vCA1 to specifically knockdown the presynaptic Nrg1 in ACC-vCA1, AAV2/9-DIO-shRNA (scramble) served as control (shNT), after one month of virus expression (3 months-old), the mice were performed to behavior tests (Figure 6D). We found that the depressive-like behaviors in 5xFAD mice were relieved by presynaptic downregulation of Nrg1 compared to the control group (Figure 6E-F). Considering the attenuation of depression of AD mice by knocking down Nrg1, we wondered whether the cognition impairments of these AD mice could be rescued. To figure that out, the same mice were subjected to water maze tests after another 3 months. Notably, knocking-down pre-synaptic Nrg1 of ACC-vCA1 significantly relieved spatial learning and memory deficits in 5xFAD mice compared with the control group (Figure 6G-J).

We then examined whether downregulation of Nrg1 affects the synaptic functions of ACC and the synaptic plasticity of vCA1 in AD mice. AAV2/9-DIO-shNrg1-eGFP or the control AAV2/9-DIO-shRNA (scramble) were injected into the ACC of Camk2a- cre and Camk2a-cre/5xFAD mice to specifically suppressed the expression of Nrg1 in Camk2 positive neuron in control or AD mice. The whole-cell patch recording of eGFP-labeled cells in the ACC was performed (Figure 6K). The reduction of Nrg1 was confirmed by immunofluorescent staining (Figure 6—figure supplement 2A and B), RT-PCR (Figure 6—figure supplement 2C) and western blotting (Figure 6—figure supplement 2D and E). We first observed dramatically reduction in sEPSC of excitatory neuron within ACC in 3-month-old 5xFAD mice which may be induced by early amyloid plaque deposition. Nrg1 knockdown significantly increased both amplitude and frequency of sEPSC in ACC of AD mice compared with control (Figure 6L-N). Furthermore, the paired-pulse facilitation (PPF) ratio in 5xFAD mice was significantly larger than WT mice, indicating that presynaptic function was disturbed in ACC of 3 months-old 5xFAD mice. Nrg1 knockdown also rescued this defect which further support the important pre-synaptic function of Nrg1 in AD (Figure 6O). We did not observe difference between Input (stimulation intensity) - Output (EPSC amplitude) responses in the above mice, indicating postsynaptic AMPA-mediated responses were not affected (Figure 6P). Finally, the LTP impairment observed in vCA1 of AD mice were restored by Nrg1 knockdown in ACC region (Figure 6Q-R). These data suggested that ectopic expression of Nrg1 specifically impaired presynaptic glutamate release and LTP formation in ACC-vCA1 circuit in AD mice.

We then examined whether the up-regulation of Nrg1 is sufficient to trigger the depression and cognition impairment in WT mice. Cre -dependent AAV-Nrg1-mCherry (AAV-OENrg1) or the control AAV-mCherry virous were injected into the bilateral ACC of Camk2α-Cre mice. The overexpression of Nrg1 was confirmed by western blotting (Figure 7A-C). Notably, we found that the ACC Nrg1 overexpressing mice exhibited depressive behaviors as tested by TST and FST (Figure 7D). In addition, extra-expression of Nrg1 by injection of AAV-Nrg1 in the ACC abolished the anti-depression effect (Figure 7E-G) and cognition improvement effect (Figure 7H-K and Figure 7—figure supplement 1) of ACC-vCA1 chemogenetic activation in 3-months-old and 6-month-old AD mice. Taken together, these results describe that the ACC^Nrg1^-vCA1 circuit in AD mice links depression and memory defects in AD mice.

**Figure 7.**
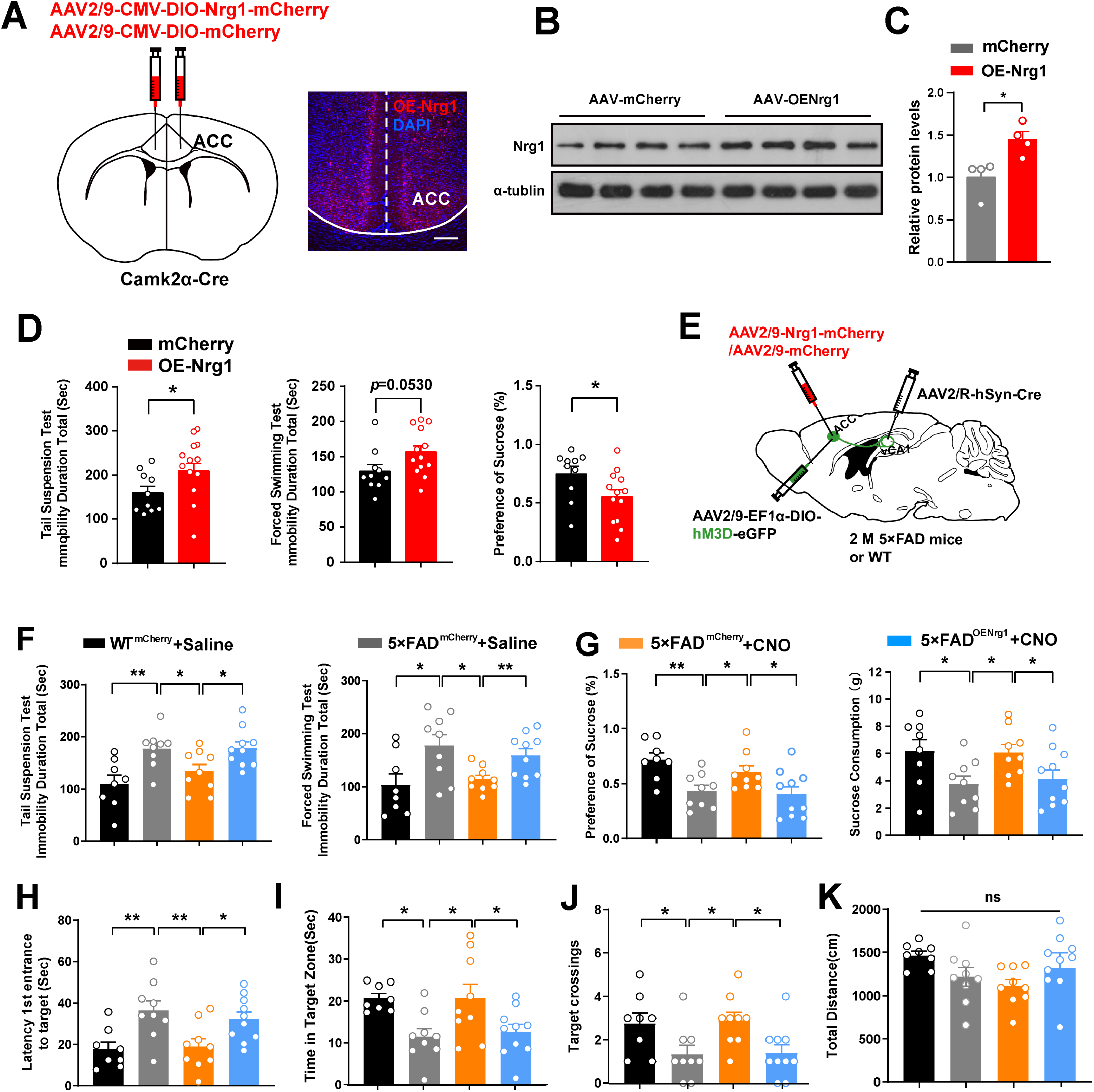
Activation of ACC^Nrg1^–vCA1 circuit ameliorates depression in early stage of 5xFAD mice. (A) The schematic diagram and representative image of the overexpressing virus AAV-CMV-DIO-Nrg1 was bilateral injected into the ACC of WT mice (scale bar 100 μm). (B, C) Western blots showing efficient Nrg1 overexpression in the ACC. unpaired t-test, n = 4 mice per group. (D) Depression-like behaviors in the TST, FST and SPT induced by over-expressing Nrg1 in ACC. (n = 8/9/9/10 mice per group). (E) Schematic of ACC–vCA1 chemogenetic activation and simultaneously ACC Nrg1 overexpression virus injections in 2 months-old WT or 5xFAD mice. (F) In the TST and FST, 5xFAD-OENrg1 with CNO treatment mice show more immobility time. (G) In the SPT, the chemogenetic activation of ACC-vCA1 inputs increased sucrose preference and consumption in 5xFAD-mCherry mice but not 5xFAD-OENrg1 mice. (H, K) In the MWM, chemogenetic activation of ACC–vCA1 inputs increased the time required from entrance to the target platform (H), target time (I) and number of target crossings (J) in 5xFAD-mCherry but not 5xFAD-OENrg1 mice without changing distance travelled (K). Data represent mean ± SEM. One-way ANOVA analysis, ns: not significant, *P < 0.05; **P < 0.01. Figure 7**—source data 1.** Source data indicating upregulation of Nrg1 in ACC induced depressive symptoms and abolished the antidepression effect and cognition improvement effect of ACC-vCA1 chemogenetic activation in AD mice. Figure 7 **— figure supplement 1.** Activation of ACC^Nrg1^–vCA1 circuit rescues depression-associated memory impairment in 5xFAD mice.

## Discussion

In the present study, a novel circuit and synaptic pathway mechanism underlying depression in the early stage of AD was reported. The chemogenetic activation of the ACC-vCA1 neural pathway was found to restore glutamatergic synaptic transmission and postsynaptic plasticity in 5xFAD mice. Besides, the effects reversed depressive-like behaviors and spatial memory deficits. Such changes were mediated by Nrg1 downregulation, and knocking down the expression of Nrg1 prevents pre-synaptic deficits in ACC of AD mice. As a result, these results suggest the restoration of the synaptic function in the ACC^Nrg1^-vCA1 circuit as the therapeutic target in early AD with depression.

Given the importance of both ACC and vCA1 in cognition and emotion, and in major brain areas receiving projections from ACC, it can be seen that vCA1 is a limbic brain region mainly included in AD’s emotional-affective spatial memory (Bian et al., 2019; Pi et al., 2020; Rolls, 2019). Since ACC and vHPC are both engaged early during the memory encoding process and are implicated in the processing of emotional-associated memory (Sacchet et al., 2017), it was speculated that the ACC-vCA1 pathway has a close association with the occurrence and development of depressive-associated memory deficits of dementia. In the research on human brain neuroimaging, it is found ACC is overinhibited in depression (Hamani et al., 2011; Lane et al., 1998), which supports the view that ACC plays a crucial role in depression symptoms. There are a lot of studies on the linkage between ACC abnormality and depression in AD. For instance, the research on PET and SPECT showed that there are abnormal blood flow, neural activity, and ACC metabolites in AD patients with depression (Xiaozheng Liu et al., 2017). The ventral hippocampal CA1 has a temporary role in the retrieval of memories, and they are stored in neocortex regions like ACC permanently (Alvarez, Zola-Morgan, & Squire, 1995; J. J. Kim & Fanselow, 1992). Furthermore, a specific role for ACC in learning and memory at a remote time was also found in the additional and more accurate experiments that adopt chemogenetic inhibition of glutamatergic neurons in ACC or their projections to vHPC (J. Kim, Wasserman, Castro, & Freeman, 2016; Weible, 2013).

It is common to see depression in pre-clinical AD, which is likely to show the early manifestation of this disease before the appearance of cognitive impairments (Visser, Verhey, Ponds, Kester, & Jolles, 2000). From that aspect, much evidence shows that depression is strongly related to AD. As a result, that mental illness is considered to be the risk factor for AD or the prodromic AD phase (Modrego & Ferrández, 2004). Accumulating evidence proved that depression is an early feature of AD (Dreimüller et al., 2019; Geerlings et al., 2000). At the same time, amyloid deposition leads to disrupted functional connectivity which impairs networks that are implicated in depression (Mahgoub & Alexopoulos, 2016). It was found that younger AD mice without cognition defects exhibit depressive-like behaviors (Figure 2). Synaptic impairment is considered a core feature of AD and depression. As the most abundant excitatory neurotransmitter for cortical and hippocampal pyramidal neurons, glutamate plays a significant role in both memory and cognition (Francis, Sims, Procter, & Bowen, 1993). We found that ACC glutamatergic transmission to vCA1 is suppressed in the early stage of AD mice. At the same time, chemogenetic or photogenic activation of ACC-vCA1 synaptic transmission could ameliorate depressive and cognitive defects in AD mice (Figure 2 and Figure 4). The accumulation of β-amyloid deposition in ACC at the early AD stage is helpful to AD-induced depression, which does not affect cognitive function during this stage.

In the research findings, it is indicated that ACC–vCA1 inputs play a critical role in associating depression with memory impairment in AD. The neuronal dysfunction of ACC mediates the impaired project to vCA1. To figure out the underlying mechanism for ACC impacted synaptic function, we performed RNA-seq in 3 months-old 5xFAD mice and identify Nrg1 as one of the key regulators. Nrg1 refers to a transmembrane protein containing an intracellular cytoplasmic domain that via binding of its extracellular domains to ErbB4 tyrosine kinase receptors, contributes to neurotransmitter release (Jin et al., 2011; Talmage, 2008). This is considered a risk gene for depression identified by a recent GWAS study (Howard et al., 2019; Schork et al., 2019) and MAGMA (Multimarker Analysis of GenoMic Annotation) analysis. Furthermore, depressed patients show a disruption of Nrg1 signaling (Mahar et al., 2017; Mahar et al., 2011). This research indicates Nrg1 expression could increase in ACC pyramidal neurons which can be labeled with the projective neurons from vCA1 in 5xFAD mice that are 3 months old (Figure 6). The high level of Nrg1 in the ACC-vCA1 circuit has a positive correction to the manifestations of early depression and cognitive disorders. Nrg1 suppression restored synaptic deficits in the ACC region and rescued early depressive-like behaviors and subsequent cognitive impairment in AD mice. Our data also found that overload of Nrg1 in ACC strongly abolished the anti-depressive effect of ACC-vCA1 chemogenetic activation in 5xFAD mice, which indicates Nrg1 plays a crucial role in encoding assessment information that is related to negative emotion and memory. As a result, Nrg1 levels are considered to decide ACC^Nrg1^-vCA1 circuit-associated amelioration of depression and memory deficits in AD mice. Taking into account no efficient therapy for AD at present while depression symptoms are amenable to treatment, these research results are likely to assist in informing risk stratification and perfecting treatment for AD patients with depression and dementia.

The study has a few limitations. First of all, we only examined the monosynaptic projection from ACC to vCA1, while ACC and vCA1 share reciprocal projections, and CA1-ACC encodes contextual information in fear (Liu et al., 2022). As a result, there needs to have further analysis on if the anti-depressive information can be transmitted back to ACC for the final integration in an ACC-vCA1 loop circuit. Secondly, one of the synaptic mechanisms of Nrg1 in the ACC-vCA1 pathway is only tested. However, we also found that several genes exhibit increased expression in ACC of 3-month-old AD mice by RNA-seq analysis. Thus, there is likely to be another synaptic molecule changed in ACC-vCA1 of AD mice in addition to Nrg1, which remains to be further tested in the future. Thirdly, it might be worth to further assessing whether the antidepressant drugs, e.g., ketamine, can exert neuroprotective effects against AD by using the rescue ACC^Nrg1^–vCA1 neural pathway.

To sum up, our research results of the ACC-vCA1 pathway revealed a relationship between depression and AD at the level of a neural circuit. The ACC-vCA1 circuit can be targeted as a meaningful method to ameliorate depression in AD and is beneficial for AD prevention.

## Materials and methods

### Key resources table

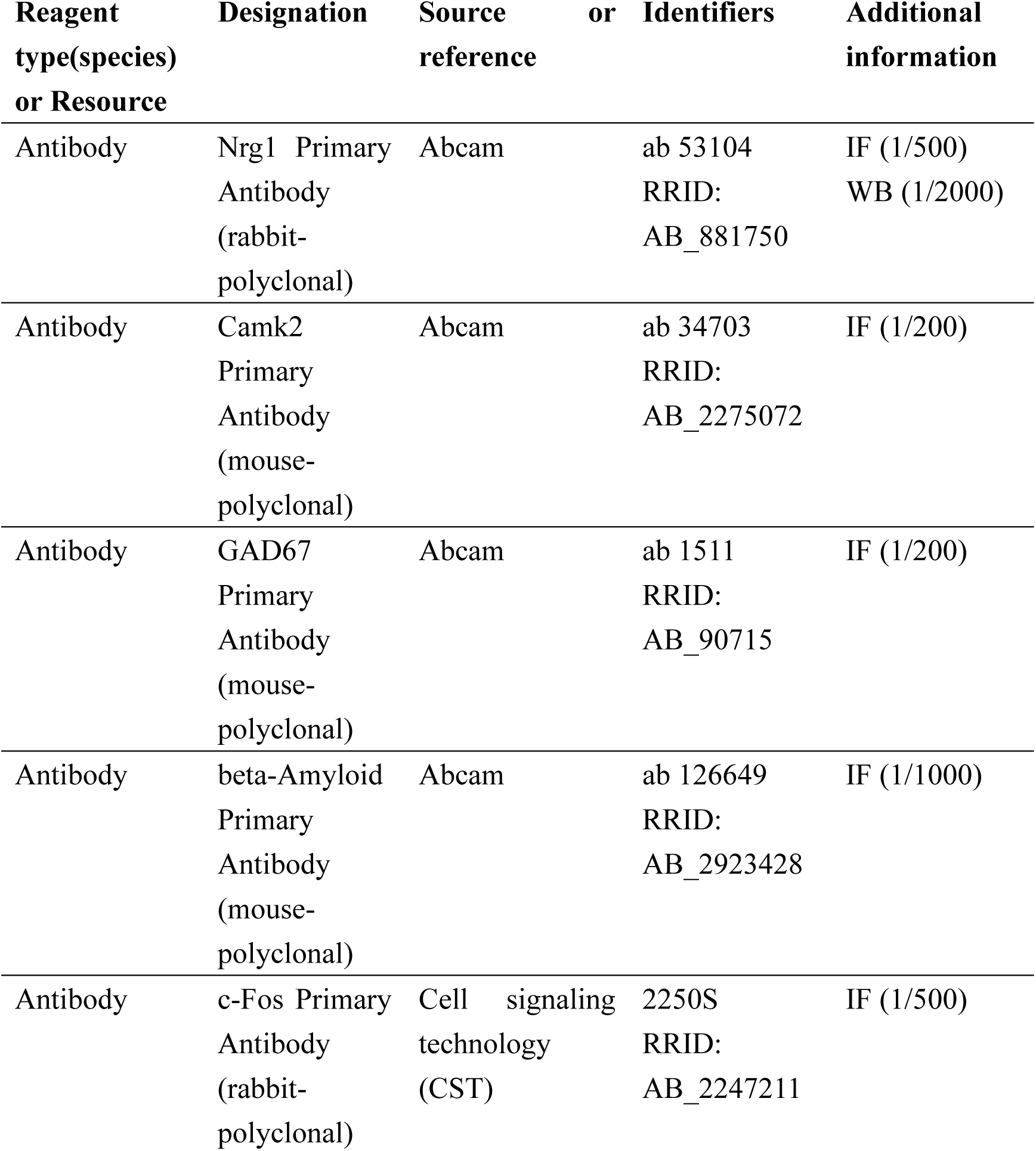
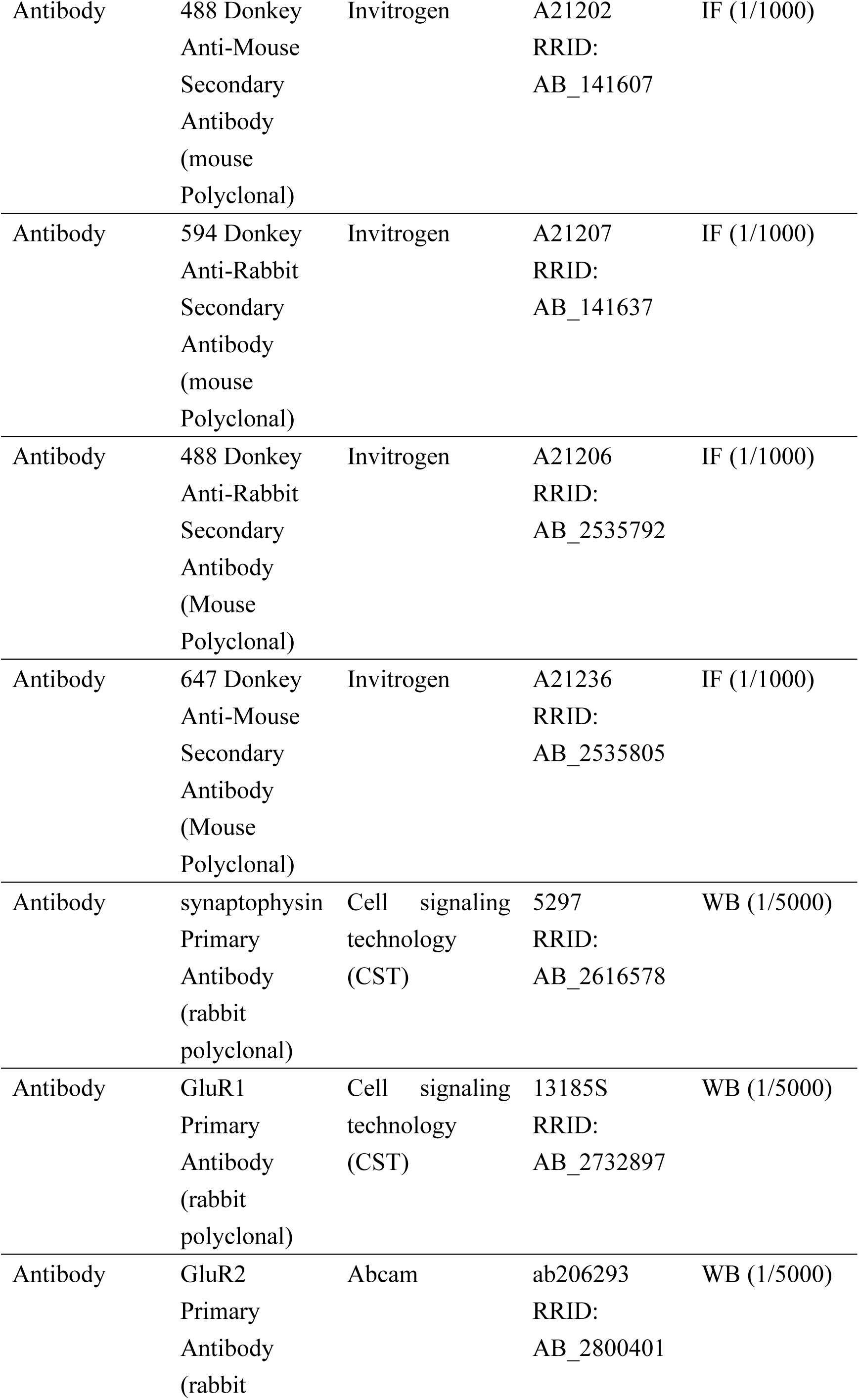
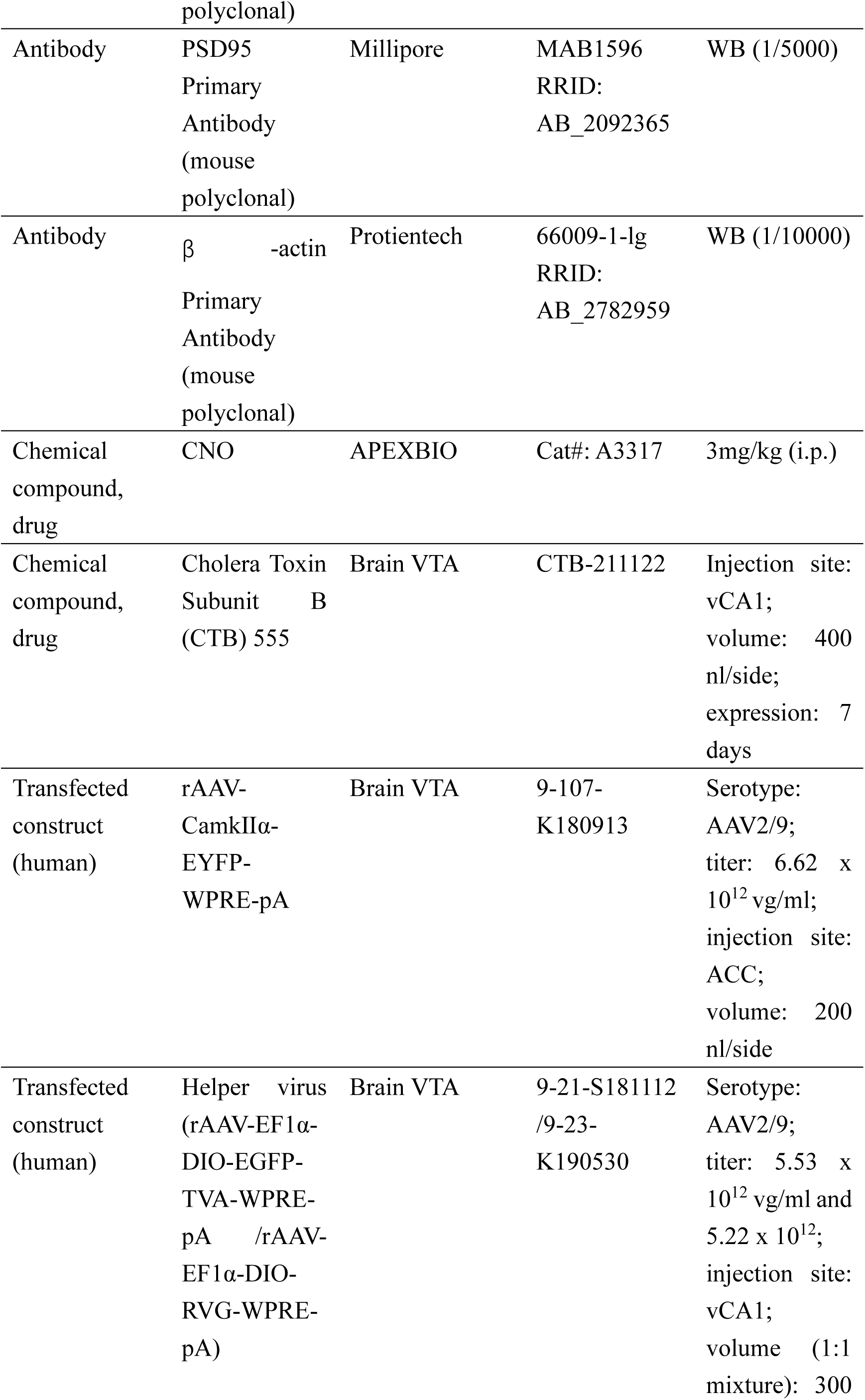
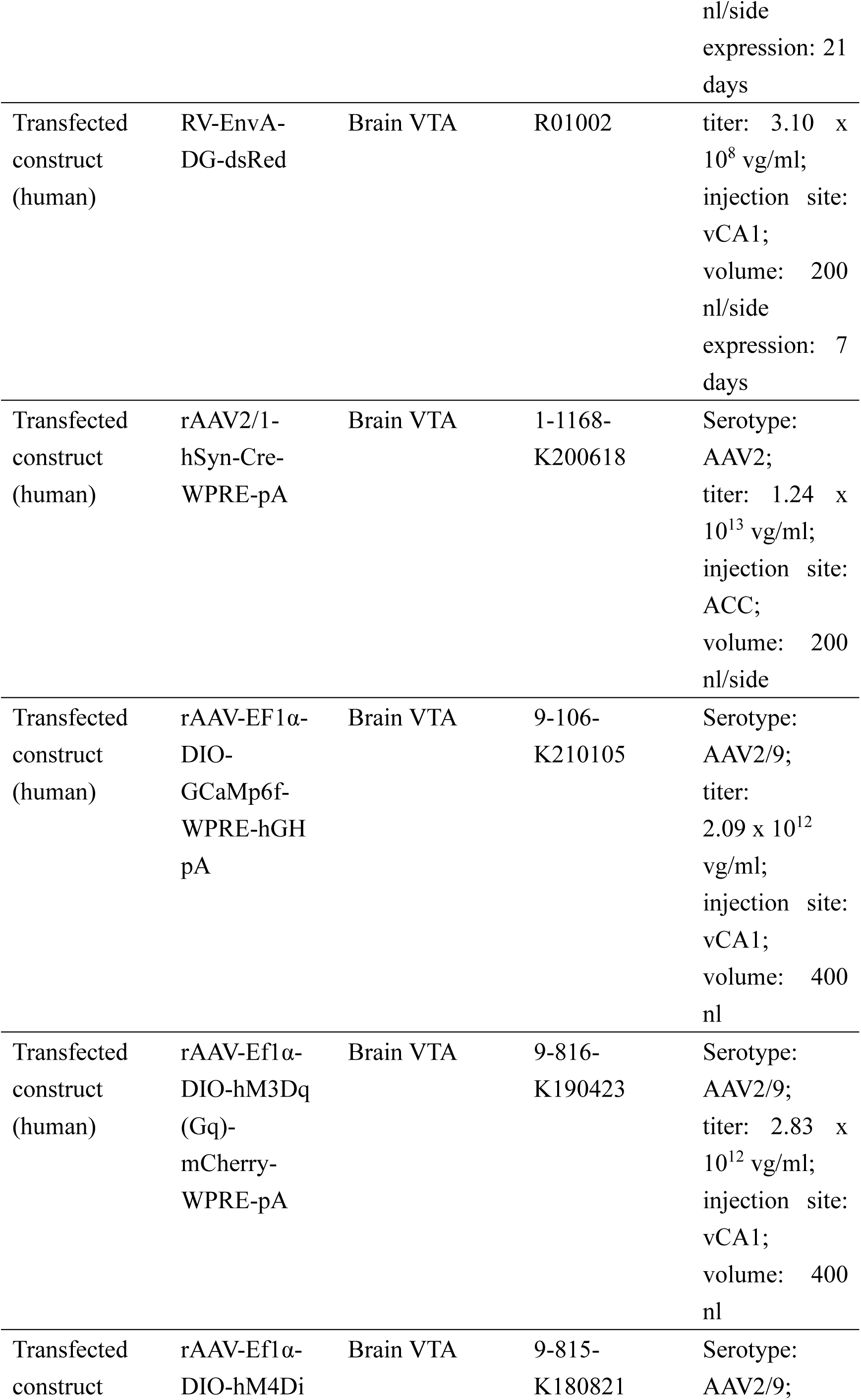
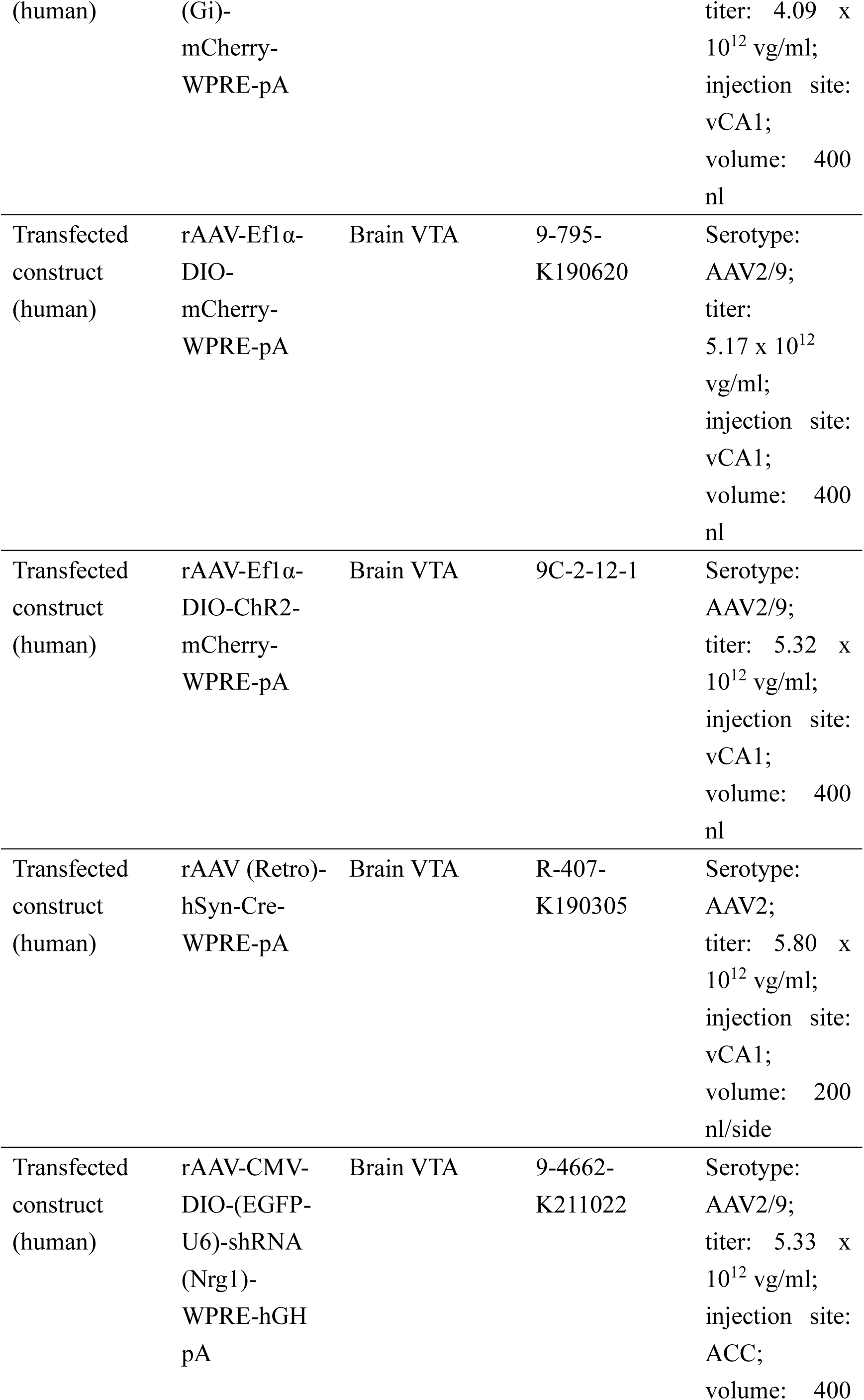
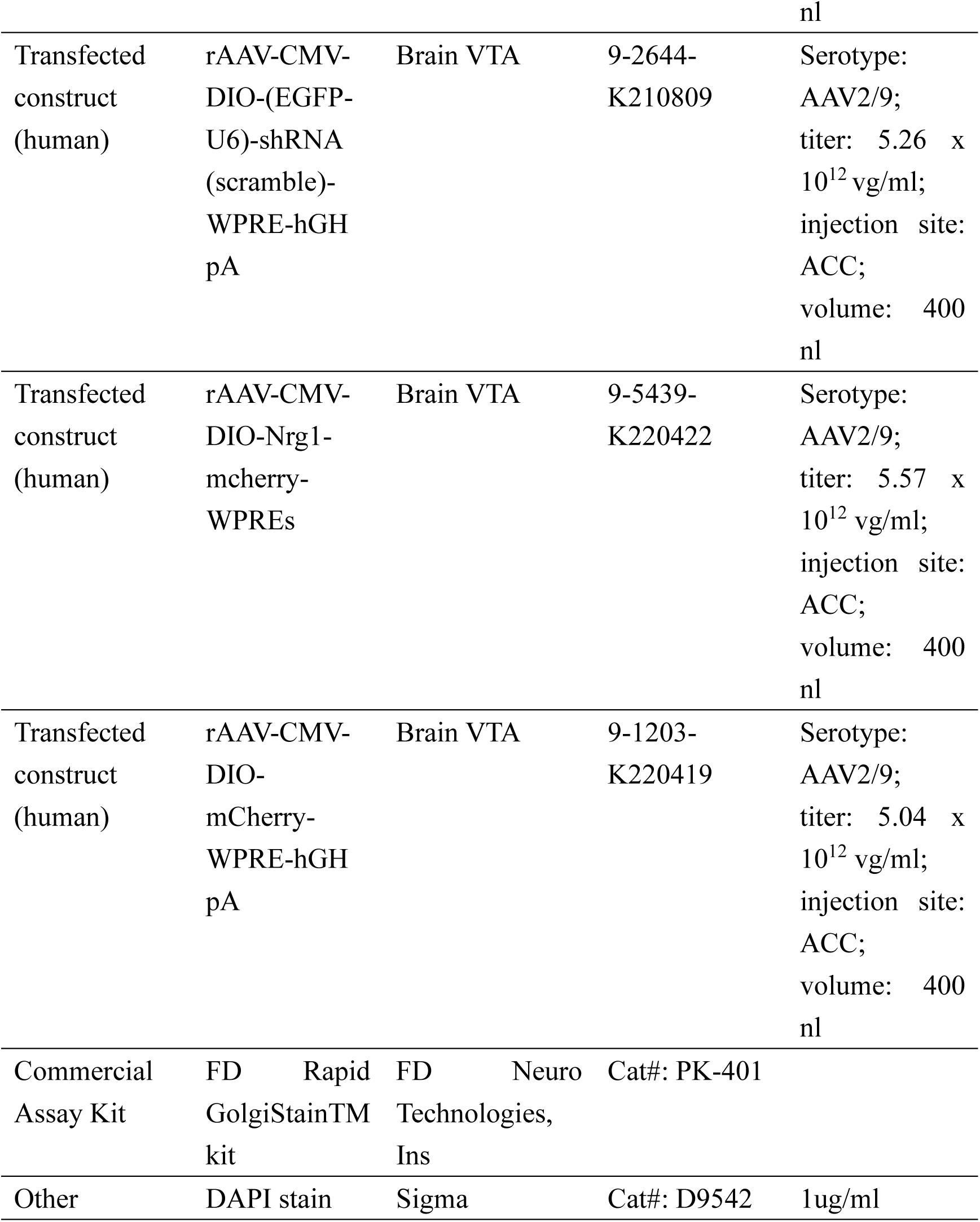

### Animals

Male 5xFAD mice (p40-55) and age-matched control mice were purchased from Model Animal Research Center of Nanjing, 3-, 6-month age and littermate mice were used for behavioral tests. Camk2α-Cre transgenic mice were purchased from Jackson Laboratory (USA). All mice were group-housed in plastic cages under SPF conditions with disposable bedding on a standard 12/12h light and dark cycle with freedom of food and water.

### Brain stereotaxic surgery and virus microinjection

The kind of virus, the time of expression and injection volume are listed in the Key Resources table. For stereotaxic surgery, animals were deeply anesthetized with intraperitoneal injection of sodium pentobarbital (35 mg/kg) and immobilized in stereotaxic frame. Then a glass micropipette attached to microsyringe (1μl, Hamilton, NV, USA) were used for virus delivery at flow rate of 20nL/min. The body temperature was still kept at 36°C throughout the procedure using a heating mat. After each microinjection, the microsyringe remained in the injection site for 10 min at the conclusion of infusion to avoid virus overflow. The virus injection bregma coordinates were as follow: ACC (AP: 1.1 mm, ML: ± 0.38 mm - ± 0.42 mm, DV: -2.1 mm), vCA1 (AP: 3.3 mm, ML: ± 4.1 mm - ± 4.5 mm, DV: -3.5 mm).

For whole brain quantitative mapping, a viral-based anterograde tracer, rAAV- CamkⅡα-EYFP was injected into the ACC of 3-month-old 5xFAD and WT mice respectively. The mice were allowed to survive for four weeks.

For retrograde transsynaptic tracing, the recombinant RV system was injected into the right side of the vCA1. The RV, whose glycoprotein (G) gene is deleted from the genome (RV-EnvA-DG), cannot spread across synapses. G complementation enables the transsynaptic spread of RV-EnvA-DG to presynaptic neurons. The virus particles formed by the RV were packed with the fusion protein of the envelope protein (EnvA) of the recombinant avian sarcoma virus, which can specifically recognize the EnvA receptor TVA to specifically infect cells. With the aid of helper viruses, RV-EnvA-DG can be used to perform cell-type-specific retrograde transsynaptic tracing in specific types of neurons.

For ACC Nrg1 knockdown intervention, the short hairpin RNA (shRNA) for Nrg1 was designed using online BLOCK-iT™ RNAi Designer program (Invitrogen) and the shRNA sequences targeting neuregulin were: 5’-GCCTCAACTCCGAGAAGATCT-3’. For the presynaptic Nrg1 knockdown experiments, rAAV- CMV-DIO-shRNA- -eGFP or rAAV-CMV-DIO-shRNA(scramble)-eGFP was delivered into the ACC, and AAV2/Retro-hSyn-Cre was injected into the vCA1 in 5xFAD mice.

For Nrg1 overexpression (OE-Nrg1) experiments, overexpressed viruses were bilaterally delivered into the ACC of Camk2-Cre mice, and subsequently allowed to recover for 4 weeks prior to undergoing behavioral testing.

### Behavioral assay

In all behavioral tests, mice were assigned randomly to the various experimental groups without restricted randomization. Investigators conducting the experiments were blinded to the allocation sequence, and we used the least number of animals in our research to prove the assumptions according to our preliminary experiments or references. Some subjects show little or no movements, i.e., they just sited there during the test, and we think these individuals did not show normal activity and motivation, therefore not appropriate to include their data in the following analysis (Figure 2 and Figure 4).

### Tail suspension test (TST)/Forced swimming test (FST)

TST and FST were both performed to measure the despair behavior. For TST, mouse tail was separately hung about 50-cm above the table with the help of tape coated with adhesive. For FST, mice were individually placed in a hyaline cylinder (15 x 25 cm) filled with clean water (maintained at 22 ± 2°C) to a depth of 15 cm. Water in the cylinder was changed frequently to prevent behavioral alteration among animals due to used water. During the 7-min process (1 min for adaptation and 6 min for recording), the experiments were monitored from the side with a video tracking system (Smart 3.0). The immobility time in individual tests was recorded. Mice in TST were considered immobile when they were passively suspended/and completely motionless, and immobility also included passive swaying. The immobility in FST was defined as the only motions which the animal’s head above the water without any other movements. The mice were performed another twice independent repeated behavioral experiment for one month later.

### Sucrose preference test (SPT)

SPT was performed to measure anhedonia behavior, which is a core symptom of depression. The SPT consists of the adaptive and testing phase. The first phase, mice were acclimatized to two bottles of 1% sucrose solution (w/v) in each cage. 24 h later, one of the bottles was altered to water for another 24 h. The positions of the two bottles were switched after 12 hours in order to avoid side bias. The second phase is that mice were deprived of water and food for 24 h and housed individually in the cages with free access to two bottles with one containing water and another sucrose solution for 12 h, the total volumes of consumed sucrose solution and water were taken note, and sucrose preference (SP) was also calculated. SP (%) = sucrose consumption / (sum of the sucrose consumption and water consumption) x 100%.

### Morris water maze test (MWM)

A tank 120 cm in diameter filled with fresh water (22 ± 2°C) was made opaque with a white, nontoxic titanium dioxide. The walls of tank were labelled with several distinct extra-maze cues. The first day they were training by a circular platform. If a mouse did not find the platform within 1 min, it was guided to the platform without struggling and allowed to remain for 10 s. The platform location remained the unchanged throughout the training, but the start points (north, south, east or west) varied randomly during the daily training. Mice received four trails per day for 4-6 consecutive days with a 15 min intertrial interval until the mice found platform within 1 min. The maximum time allowed per trial was 60 s. Escape latency was recorded for each trial for spatial memory acquisition. On day 7, the platform was removed and the mice were gently placed in the NE direction swimming for 1 min. The percentage time spent in every of the four quadrants and the number of target (platform) area crossings, mean speed, total distance was recorded. For optogenetic activation, photostimulation was delivered onto vCA1 neurons 1 min before training and testing every day.

### Immunofluorescence staining assay

The mice were deeply anesthetized flowing by perfused transcardially with ice-cold 0.1 M phosphate-buffered saline (PBS) shortly followed by 4% paraformaldehyde (PFA). with PBS and PFA. The brains were subsequently removed and post-fixed in PFA at 4°C overnight and immersed in 30% (w/v) sucrose solution (4°C) for at least 48 h. Then, coronal sections (30 μm) were cut on a cryostat (Leica CM1950) and used for immunofluorescence. Brain sections were incubated in 0.3% (v/v) Triton X-100 for 0.5 h, blocked with 5% goat serum at room temperature (RT) for 1 h, and incubated with primary antibodies at 4°C for 24 h, which were visualized with the corresponding fluorophore-conjugated secondary antibodies for 2 h at RT. Then, 4,6-diamidino-2- phenylindole (DAPI) were used at the last stage. Finally, the staining sections were finally scanned and imaged using Olympus FV1000MPE-B.

For whole brain quantitative mapping, brain tissues were performed the 25 μm thickness frozen sections at eight levels of bregma 2.2 mm, 0.74 mm, -0.34 mm, -1.70 mm, -2.80 mm, -3.40 mm, -4.72 mm, and -5.40 mm. Brain section containing EYFP-positive signals and then assigned to specific brain regions based on the Allen Brain Atlas. The fraction of axons resulting from ACC neurons was determined via measuring and averaging the pixel densities of the image in each brain area. EYFP signals were then normalized in each brain region by EYFP signals of brain region injected in the same brain (Schwarz et al., 2015) and were detected with a fully automated slice scanner (VERSA 200, Leica, Germany).

### Fiber photometry in the TST

The change in neuronal activity in the TST was assayed by recording GCaMP fluorescent signals with an optical fiber recording system (RWD Life Science Co., Ltd.) as previously reported (Zan et al., 2021). In brief, an optical ceramic needle was inserted towards the vCA1 through the craniotomy after the corresponding virus injection according to Key Resources Table The mice were housed individually for 7 days for recovery. A 488 nm laser (0.01–0.02 mW) was delivered using an optical fiber recording system, and fluorescent signals were recorded. For data analysis, the original signal is demodulated and converted to Δ F/F. Next, the demodulated signal is converted to Δ F/F using an average of 30s before and after each data point as F, normalising each data point F n with the formula (F n − F)/F. We set the time point at which the mouse was going to immobile as 0. Four to seven bouts of a behavior were collected during the test. If a specific behavior occurred no more than four times within 1h, the record was replicated on the following day.

### In vivo optogenetic/ pharmacogenetic manipulations

For optogenetic activation, approximately three weeks after virus expression, an optical fiber (diameter of 200 μm, NA: 0.37, RWD Life Science Co., Ltd.) was unilaterally implanted in vCA1 and fixed using dental cement. Implantable fibers were connected to a laser stimulator (RWD Life Science Co., Ltd.) and delivered a 5-min pulse of blue light (473 nm, 2–5 mW, 20 Hz). The exact same stimulus protocol was applied for the control group. The implanted site of the fibers was checked in all mice at the end of the experiments, and row data obtained from mice in which the fibers were implanted outside the vCA1 were discarded. The output parameters were 5 ms, 20 Hz, 3 mins off, 3 mins on, ∼2 mW for behavioral experiments, which were selected and slightly modified based on previous research (Yang et al., 2016).

In pharmacogenetic studies, CNO (3 mg/kg, i.p.) was dissolved in saline given 1 hour before the beginning of the behavioral testing.

### Golgi-Cox staining

Golgi-Cox immunohistochemistry staining was performed using the FD Rapid GolgiStainTM kit, which is used for studying the dendritic spine morphology generally. First, freshly dissected blocks containing ACC and vCA1 regions were immersed in the mixture (1:1) solutions (A and B), and kept at RT for 2 weeks in the dark. Then brain tissues were put into Solution C and replaced after at least 48 hours at 4°C in the dark. Vibratome (Leica 1200S) was used to cut 200 μm thick sections. Brain slices were then histochemical stained with solutions D and E, mounted on a gelatin-coated glass slides, allowed to air dry, dehydrated and covers lipped. Golgi-stained neurons and dendritic segments from ACC and vCA1 were examined and imaged under light microscopy (AHBT3; Olympus, Tokyo, Japan). The density of neuronal dendritic length and spines was statistically analyzed using Image J (Fuji) software.

### Western blotting

The vCA1 and ACC were serially micro-dissected from 400 μm thick coronal sections. The mouse brain samples were homogenized in cold lysis buffer (RIPA, Boster China) containing a cocktail of protease inhibitors for at least 40 min, then were centrifuged at 12 000 rpm for 10 min at 4°C. We collected the supernatants and measured total protein concentration by bicinchoninic acid (BCA) protein kit (Lifetech, China). The protein samples were separated on 8–15% SDS-PAGE gels. After that, the separated proteins were transferred to PVDF membranes (polyvinylidene difluoride), and subsequently blocked in TBST containing 5% skimmed milk for 2 h at RT. Primary antibody was incubated at 4 °C overnight. The kind of antibody is listed in Key Resources Table. On the second day, the membrane was washed three times with TBST, followed by incubation with secondary HRP-conjugated antibodies for 1 hour at RT. Finally, ECL detection reagents (Pierce) were used to visual the proteins signals on the blots. Integrated gray values of each band were quantified using Image J (Fuji) software. All experimental groups and the control was replicated at least 3 times (Figure 3F, Figure 3—figure supplement 1, Figure 5H, Figure 6—figure supplement 2D-E and Figure 7B- C).

### Quantitative real time PCR

The bilateral ACC blocks from 5xFAD (3-month) and age-match control mice were collected on dry ice. Total RNA from each sample were extracted using Trizol™ Reagent in accordance with the manufacturer’s instruction (Invitrogen). The reverse transcription reaction was performed at 37°C for 15 min, 55°C for 5 min, 98°C for 5 min using High-Capacity cDNA Reverse Transcription Kit (Toyobo, FSQ-201). Targets were amplified in triplicate by quantitative real time PCR using Step OnePlus™ Real-Time PCR System (Thermo Fisher Scientific). Samples were assayed in triplicate and β-actin was used as an internal control. The PCR primers used are showed as follows:

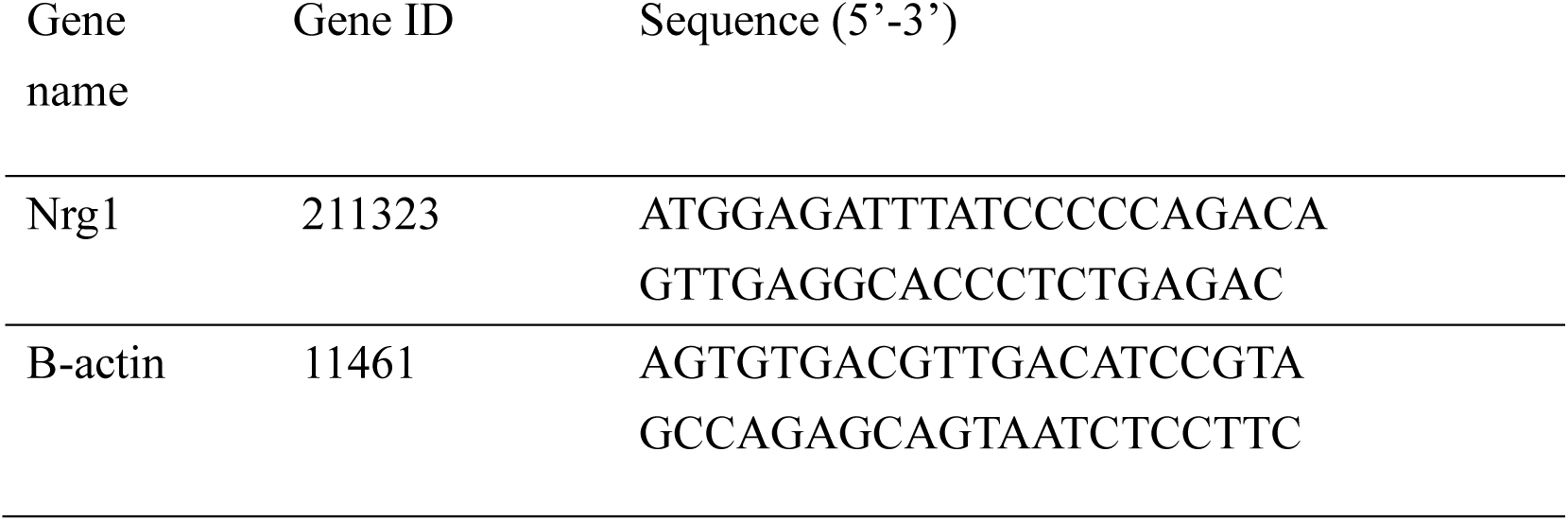

### Electrophysiology

The experimental procedures of whole-cell patch-clamp recordings were based on previous researched (Liang et al., 2020). In brief, transverse slices (400 μm thickness) containing the ACC or vCA1 were cut using Vibroslice (VT 1000S; Leica) in an iced cold solution, then subsequently were transferred to a RT-submerged chamber saturating with oxygenated (95% O_2_ and 5% CO_2_) artificial cerebrospinal fluid (ACSF) consisting of (in mM) 120 NaCl, 26 NaHCO_3_, 3.5 KCl, 1.25 NaH_2_PO_4_, 2.5 CaCl_2_, 1.3 MgSO_4_ and 10 glucoses. The recordings were performed using an amplifier (Axon Multiclamp 700B) and digitized by Clampfit 10.6 software (Molecular Devices). The patch cells were visualized using an infrared differential interference contrast microscope with a 40 × water immersion lens and an infrared-sensitive charge-coupled device camera.

#### For spontaneous EPSC and IPSC recording

CTB-555 retrogradely labeled neurons in ACC after 1 week of CTB-555 injection in vCA1 were patched. Cells were recorded at a holding potential of -70 mV for sEPSC and 0 mW for sIPSC. The potassium-based intracellular solution within the micropipettes (8-10 MΩ) contained the following (in mM): 120 K-gluconate, 5 NaCl, 0.2 EGTA, 10 HEPES, 2 MgATP, 0.1 Na_3_GTP, 1 MgCl_2_ and 10 phosphocreatine (adjusted to pH 7.2 with KOH, 290 mOsm). Neurons with a resting potential of at least -70 mV and resistance fluctuated within 10% of initial values were analyzed. Mini Analysis software (Synaptosoft, Inc) was used to analyzing the mini events. Data were filtered at 1 kHz and sampled at 10 kHz.

For paired-pulse facilitation (PPF) recording, the intervals were 35, 50, 75, 100 and 150 ms. Pyramidal neurons were current-clamped and injected 50-350 pA depolarizing currents with a step-size of 50 pA to investigate the relationship between the spike number and the intensity of injected currents. The AMPAR-EPSC were recorded at holding potentials of 50, 30, 10, 0, -20, -40 and -60 mV, The glass pipettes were solution contained (in mM): 140 CsCH3SO3, 5 mM TEA-CI, 10 mM HEPES, 1 mM EGTA, 2 mM MgCl_2_, 2.5 mM Mg-ATP, and 0.3 mM Na-GTP, and 5 mM lidocaine N- ethylchloride (QX-314) (adjusted to pH 7.3, 280 mOsm). The ratio of peak EPSC amplitudes at negative (-40 mV) and positive (50 mV) holding potentials was measured as rectification index.

#### For chemogenetic manipulations verification

Neurons expressing hM3Dq were visually identified by fluorescence of mCherry in vCA1 and recorded. Current-clamp recordings were performed to measure evoked action potentials. Spontaneous firing of action potentials (AP) was recorded in the current clamped at 70 mV. After 5 min of recording, the slices were perfused with 5 μM CNO. The total recording time for each cell was 10 min. The threshold current of the AP was defined as the minimum current to elicit an action potential with steps of 25 pA, ranging from 0 pA to 250 pA. Afterwards, the CNO was removed by washes with ASCF and cells were recorded for another 10 min.

#### For long-term potentiation recording (LTP)

Field excitatory postsynaptic potential (fEPSP) were recorded from neurons within the CA1 dendritic layer by insertion of a bipolar tungsten electrode in the Schaffer collateral pathway as the stimulating electrode. Schaffer collateral inputs to the CA1 region were stimulated with an electrode. Based on the stimulus–response curve, a stimulation intensity that evoked a fEPSP within 10% and 90% of the peak response was selected and recorded for 20 min to establish a baseline. LTP was then induced with a two-train high-frequency stimulus (5 bursts at 5 Hz, repeated 5 times at 10 s intervals, 4 pulses at 100 Hz for each burst). The field potential response following tetanic stimulation was recorded for 60 min. The LTP magnitude was quantified as the percentage change in the over 60 min after LTP induction. To verify chemegenetic activation, CNO (5μM) was perfused at 3 mL/min without recording for 30 min, then a single response was measured until the recording was stopped. For comparison of LTP magnitude, the average value of the last 10 min recordings was compared statistically.

### RNA-sequencing and analysis

Fresh ACC tissue from four 3-month-old 5xFAD mice and four litters of control mice were collected in each of the two experimental replicates. RNA was isolated and used for RNA-sequencing analysis. The quantitative analysis was performed by Guangzhou Gene Denovo Biotechnology using Xten platform. RNA-seq row data were originally filtered to obtain clean data, including removing reads with adaptors, reads with more than 10% unknown bases or low-quality reads (the percentage of low-quality bases is more than 50% in the read). High-quality clean reads were obtained by further filtering using fastp (version 0.18.0). The package software Empirical analysis of Digital Gene Expression in R (EdgeR, Gene Denovo Biotechnology Xten platform, Guangzhou, China) was used for differential expression analyses. Identify differential expressing genes (DEGs) between two different samples, performed clustering analysis and functional annotation. Genes with a false discovery rate (FDR) ≤ 0.05 and fold change ≥ 1.5 were considered to be statistically significant. GO functional enrichment and Kyoto Encyclopedia of Genes and Genomes (KEGG) pathway analysis were carried out by Goatools. The differential expression genes (DEGs) were presented in Figure 5—figure supplement-1 source data 1.

### Statistical analyses

Statistical comparisons were performed with GraphPad Prism 8.3.0 software, via one- or two-way analysis of variance (ANOVA), and unpaired Student’s *t* tests were used to compare the differences between two groups. Complete statistical details are included in source data files online for Figure 1-7. Asterisks were used to indicate significance: \**P* < 0.05, \*\**P* < 0.01, and \*\*\**P* < 0.001, and data were showed as mean ± SEM.

## Acknowledgement

We thank Z. Yuan at Beijing Institute of Basic Medical Sciences for providing 5xFAD mice and Camk2α-Cre mice. We also thank for J. Huang at Fourth Military Medical University for advice on fiber photometry experiments. The authors have no biomedical financial or potential competing interests to declare.

## Ethics

This study was performed in strict accordance with the recommendations in the Guide for the Institutional Animal Care and Use Committee at Xiamen University (XMULAC20190143). All animals were kept and maintained at Xiamen University Laboratory Animal Center.

---

**Figure 1—figure supplement 1.**
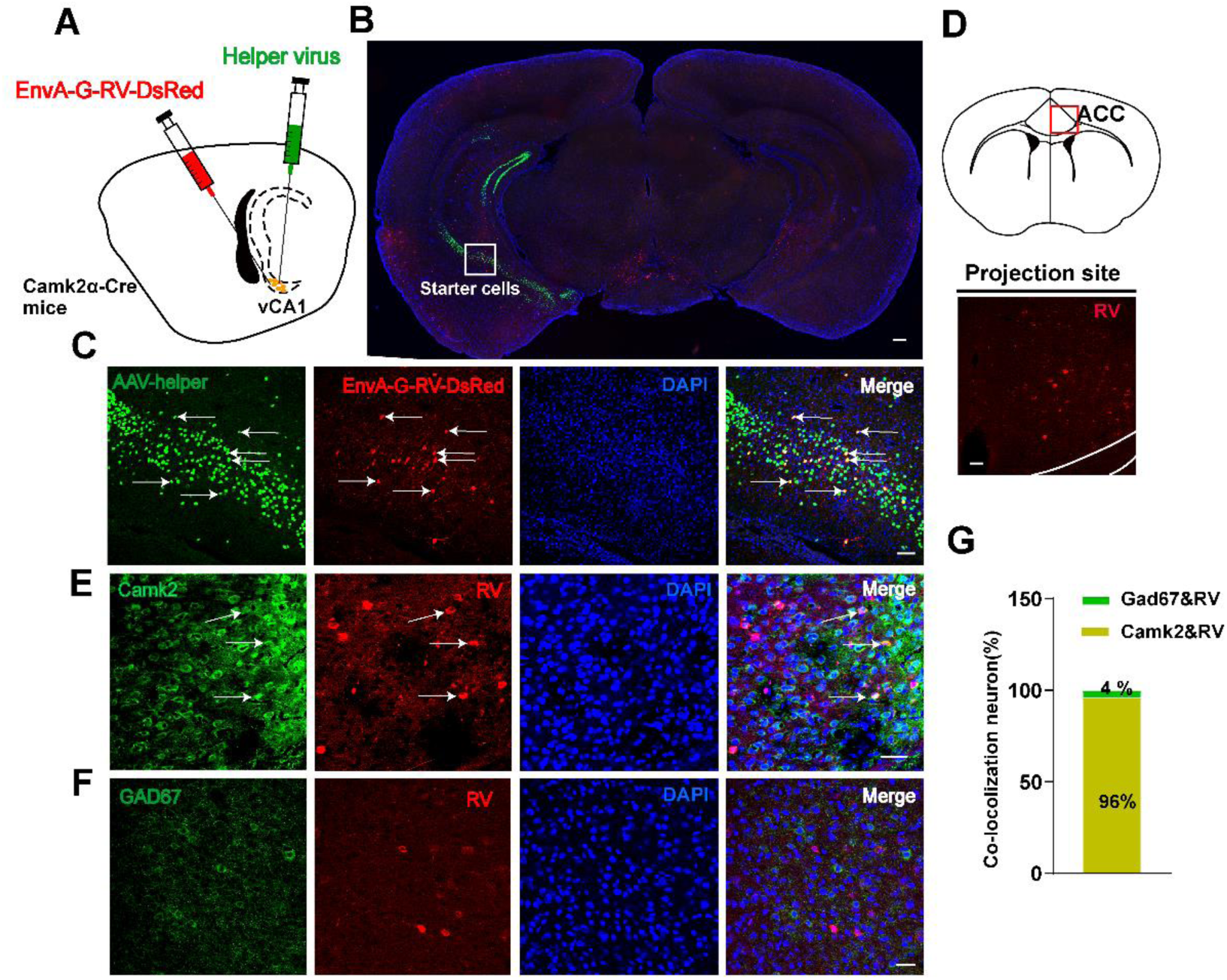
The direct glutamatergic monosynaptic projection from ACC to vCA1. Schematic diagram (**A**) and representative image (**B**) of retrograde monosynaptic tracing in Camk2α-Cre mice: the helper viruses AAV-EF1a-DIO-GT and AAV-EF1a-DIO-RVG (1:1, 200 nl, green) was injected into vCA1, and then rabies virus EnvA-pseudotyped RV-DG-dsRed (red) was also injected into vCA1. The white rectangle indicates starter cells. Scale bar = 500 μm. (**C**) The representative images of helper virus injection in vCA1. White arrowheads indicate starter cells. Scale bar = 100 μm. (**D**) The schematic diagram (top) and representative images(bottom) of RV-dsRed-labeled neurons in the ACC. red rectangle indicates the ACC region. Scale Bar = 100 μm. (**E-F**) The ACC excitatory neurons were detected by co-staining of dsRed and anti-Camk2 (**E**), and the inhibitory neurons were probed by co-staining of dsRed and anti-GAD67 (**F**) in Camk2α-Cre mice. Scale bars = 50 μm in E, 25 μm in F. (**G**) The proportion of Camk2- and GAD67-positive cells co-labeled with DsRed in ACC, respectively.

**Figure 2 — figure supplement 1.**
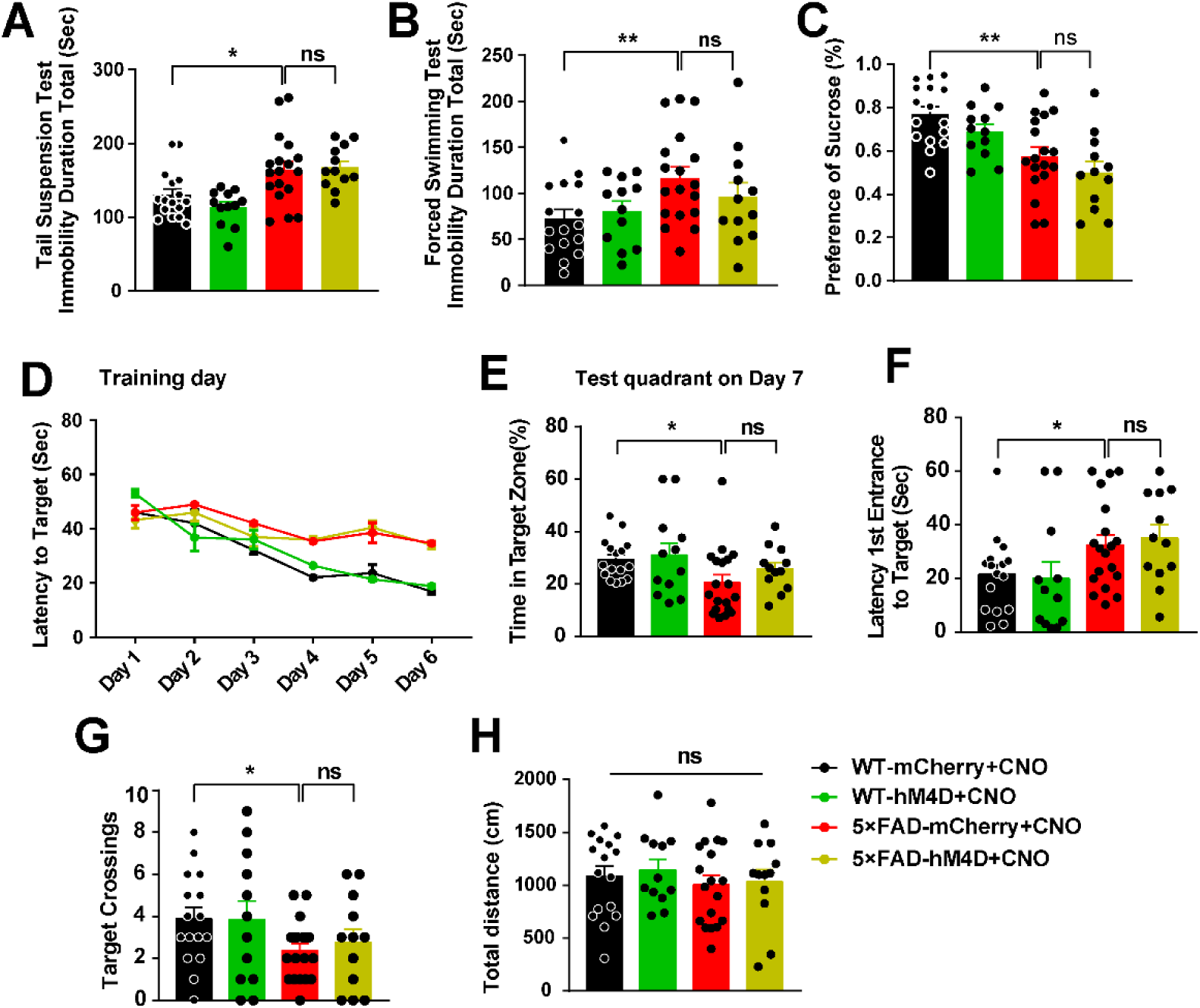
**No significant behavioral changes were found by chemogenetic inhibition of ACC-vCA1 pathway**. (**A-C**) Duration of immobility in the TST (**A**) and FST (**B**), and sucrose preference in the SPT (**C**) paradigm for 3 months-old WT and 5xFAD mice expressing hM4D and receiving CNO injections. (**D**) During MWM tests, WT or 5xFAD mice expressing hM4D and receiving CNO injections were analyzed for escape latency during a 6-day training period. (**E-G**) On the next day, mice were analyzed for time spent in the target zone (**E**), time required from entrance to the target platform (**F**) and number of target crossings (**G**). (**H**) Total distance during the training did not differ across all groups. Data represent mean ± SEM. One-way ANOVA. Mice number was n = 16 for WT-mCherry, n = 12 for WT-hM4D, n = 18 for 5xFAD- mCherry and n = 12 for 5xFAD-hM4D group, ns: not significant, \**P* < 0.05; \*\**P* < 0.01. Figure 2**—figure supplement 1—source data 1.** Source data indicating the negative effect of chemogenetic inhibition of ACC-vCA1 circuit in AD mice.

**Figure 2—figure supplement 2.**
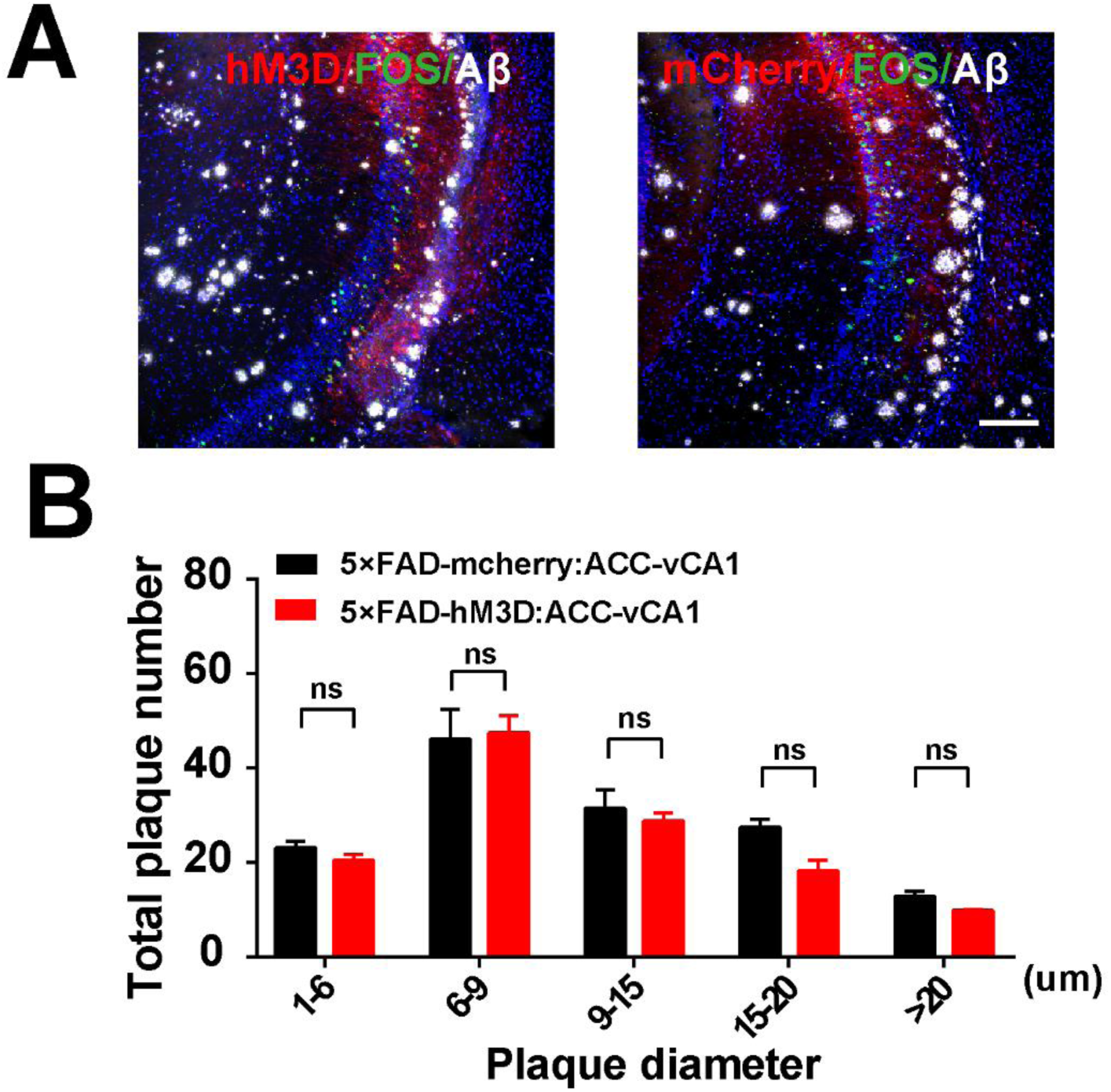
Chemogenetic activation rescues ACC-vCA1 without changing Aβ level in vCA1. (**A**) Representative co-staining of FOS/Aβ positive neurons in the vCA1 of 5xFAD mice. Scales bar = 100 μm. (**B**) Quantification of plaque number in infected cortical regions of AD mice. Data represent mean ± SEM. unpaired t-test, n = 3 mice per group, ns: not significant.

**Figure 3—figure supplement 1.**
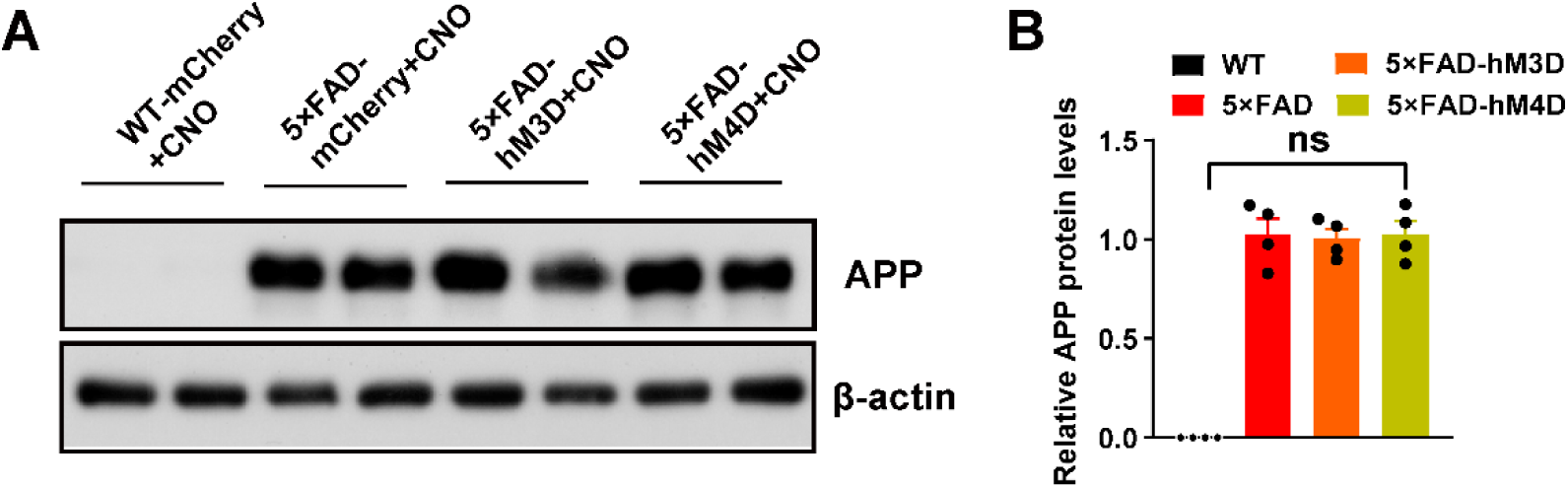
The activation of ACC-vCA1 does not affect the APP expression. (**B**) Quantification of APP in the vCA1 of 3-month-old WT and 5xFAD mice. Scale bar = 10 μm. Data represent mean ± SEM. Unpaired t-test, n = 4 mice per group, ns: not significant. Figure 3 **— figure supplement 1-source data 1.** Immunoblot and quantification of APP in hippocampal CA1 after chemogenetic activation or inhibition of the ACC-vCA1 circuit.

**Figure 5—figure supplement 1.**
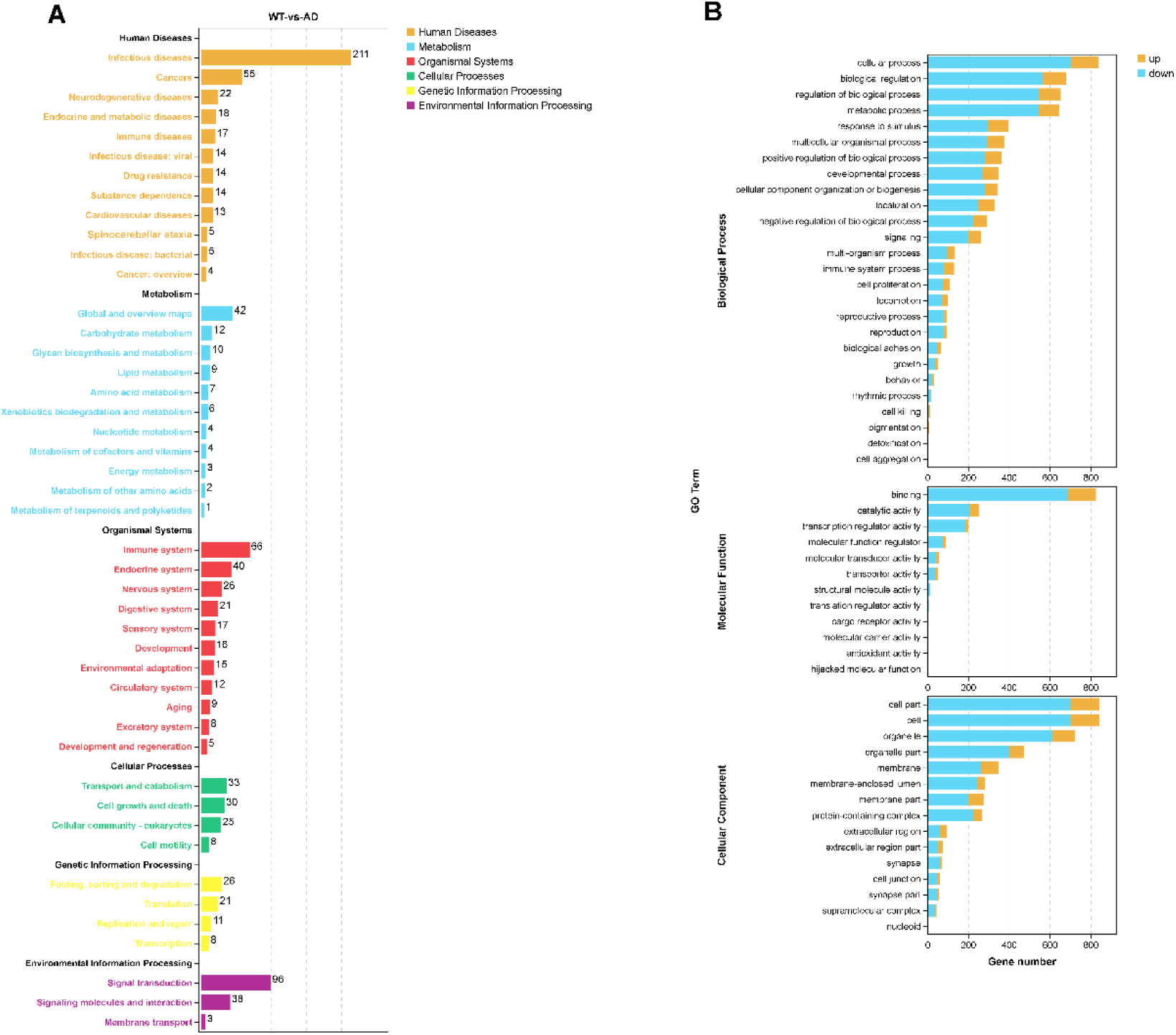
Differential expression genes-GO/KEGG analysis. (**A**) DEGs mRNAs were clustered Kyoto Encyclopedia of Genes and Genomes (KEGG) analysis. (**B**) DEGs mRNAs were clustered by Gene Ontology (GO) analysis, and the total terms of biological process, molecular function, and cellular component are shown, respectively. Figure 5**—figure supplement 1-source data 1.** Source data indicating the expression of differential genes by RNA sequence analysis.

**Figure 6 — figure supplement 1.**
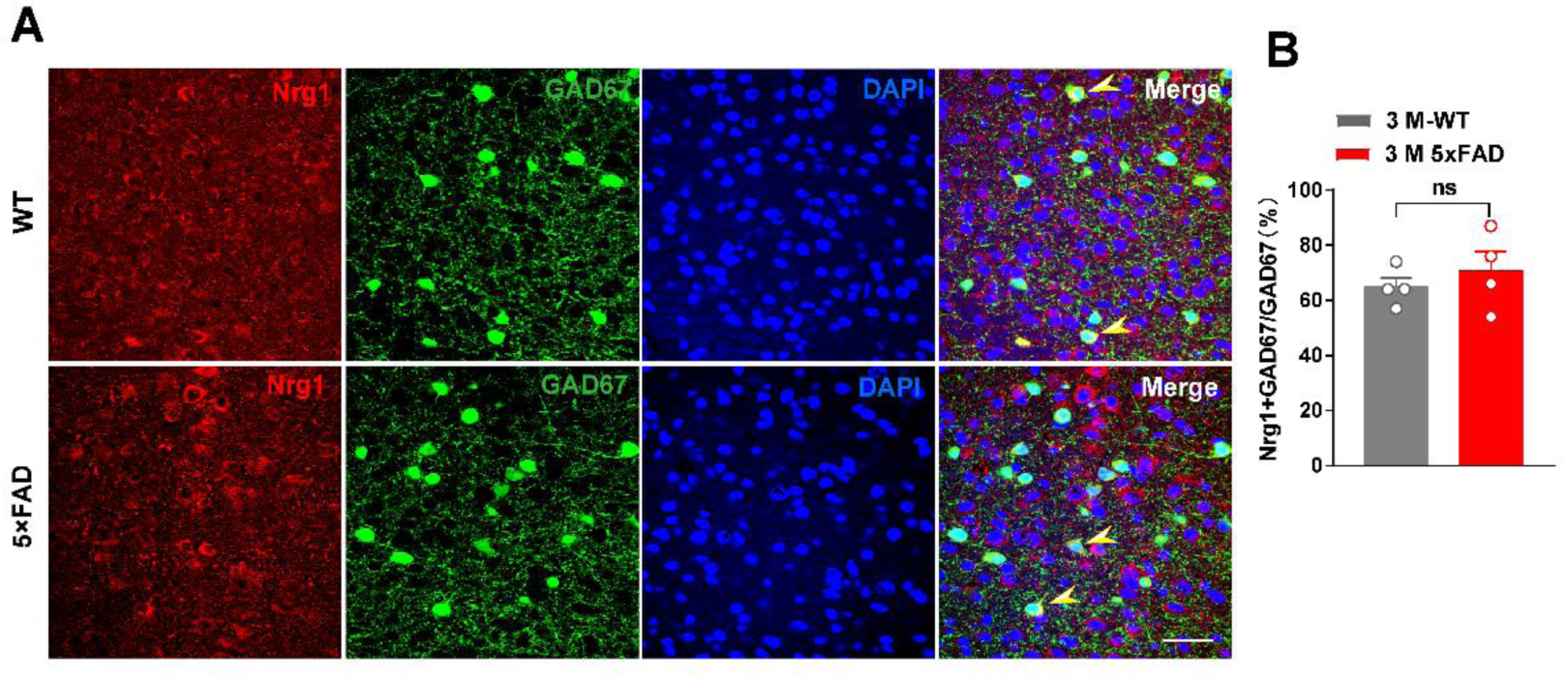
The co-existence of Nrg1 with GAD67 in ACC. (**A**) Representative immunofluorescence images showing that Nrg1 expression was found in GAD67^+^ interneurons in the ACC in both WT and 5xFAD mice. Yellow arrows indicate Nrg1/GAD co-expressed neurons. Scale bar = 25 μm. (**B**) Histogram showing that the percentage of GAD67/Nrg1- containing neurons among GAD67 containing interneurons is not different between 3 months-old WT and 5xFAD mice. Data represent mean ± SEM. Unpaired t-test, n = 4 mice per group, ns: not significant.

**Figure 6 — figure supplement 2.**
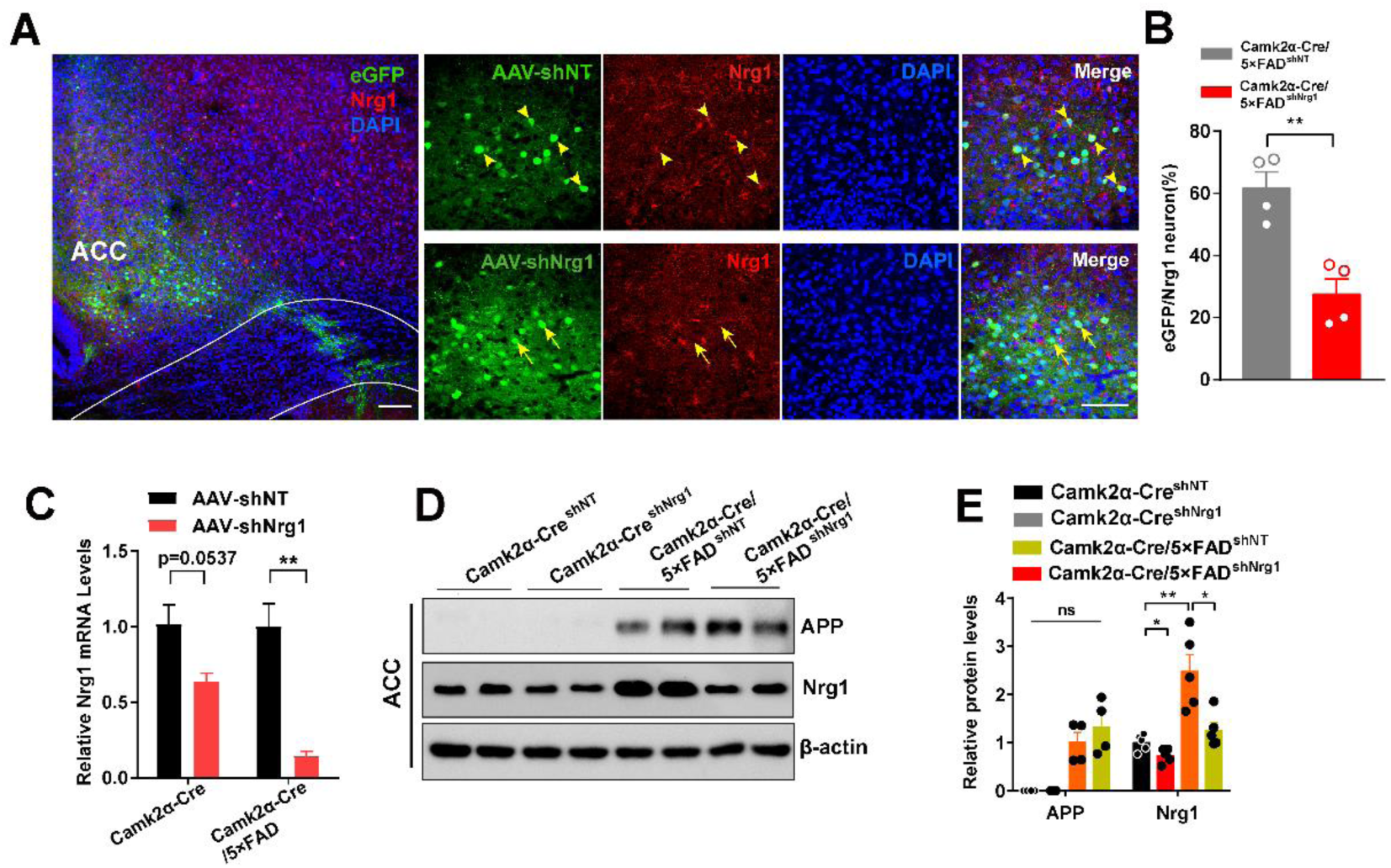
Nrg1 was knocked down efficiently by AAV-sh-Nrg1 microinjection into the ACC. (**A-B**) Double-immunofluorescence images (A) and quantification analysis (B) showing efficient Nrg1 knockdown in the ACC. Yellow arrows indicate Nrg1/eGFP double-labeled neurons, yellow arrowheads indicate eGFP-labeled neurons without Nrg1- expressing. Scale bars = 100 μm, 50 μm. (**C**) RT-PCR showing efficient Nrg1 knockdown in the ACC of Camk2α-cre and Camk2α-cre/5xFAD mice. (**D-E**) Western blots analysis showing efficient ACC Nrg1 knockdown without change in APP. Data represent mean ± SEM. Unpaired t-test, n=5 mice per group, ns: not significant, \**P* < 0.05, \*\**P* < 0.01; \*\*\*\**P* < 0.0001. Figure 6**—figure supplement 2-source data 1.** Immunoblot and quantification of APP and Nrg1 after viral knockdown Nrg1 in ACC.

**Figure 7 — figure supplement 1.**
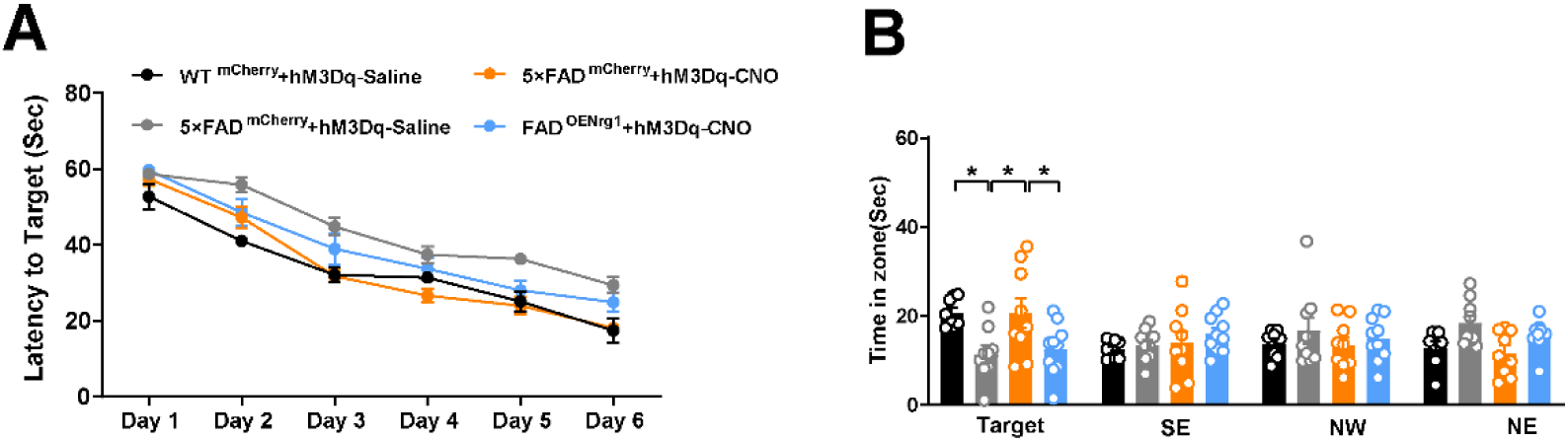
Activation of ACC^Nrg1^–vCA1 circuit rescues depression-associated memory impairment in 5xFAD mice. (**A**) During Morris water maze tests, WT/5xFAD- mCherry mice and WT/5xFAD-OE-Nrg1 mice with CNO or saline treatment were analyzed for latency of escape during a 6-day training period. (**B**) On the next day, mice were analyzed for time spent in the target zone and other three quadrants (SE, NW and NE). Data represent mean ± SEM. n = 8/9/9/10 mice per group, one-way ANOVA analysis, ns: not significant, * *P* < 0.05; compared with the control group. Figure 7**—figure supplement 1-source data 1.** Source data indicating overexpression of Nrg1 in ACC abolish the cognition improvement effect of ACC-vCA1 chemogenetic activation.

